# CA1 Engram Cell Dynamics Before and After Learning

**DOI:** 10.1101/2024.04.16.589790

**Authors:** Amy Monasterio, Caitlin Lienkaemper, Siria Coello, Ryan Senne, Gabriel K. Ocker, Benjamin B. Scott, Steve Ramirez

## Abstract

A fundamental question in neuroscience is how memory formation shapes brain activity at the level of neuronal populations. Recent studies of hippocampal ‘engram’ cells—neurons identified by learning-induced immediate early gene (IEG) expression—propose that these populations form the cellular substrate for memory. Previous experimental work suggests that cells are recruited into engrams via elevated intrinsic excitability and that learning drives coactivity among these cells to support retrieval. Despite this, an understanding of how engram dynamics evolve across learning and recall remains incomplete. Here, we combined activity-dependent genetic tagging with longitudinal two-photon calcium imaging to track CA1 engram population dynamics before and after fear conditioning. Our results reveal that engram activity is modulated by intrinsic dynamics, behavioral state, and stimulus-cued reactivation. Consistent with the idea that intrinsic dynamics bias engram allocation, spontaneous activity during rest predicted future engram membership—up to two days prior to *Fos* expression. In parallel, we found sequential activity during locomotion recruited both engram and non-engram cells, but that engram cells were less modulated by velocity. Furthermore, after fear conditioning, within the engram population, we identified a subset of cells that increased their correlations after fear learning, specifically during quiet rest. We also used a trace fear conditioning paradigm to show that conditioned stimulus presentation drove elevated activity and stable correlations amongst engram cells, demonstrating learning-dependent reactivation. Finally, computational modeling of CA3-CA1 circuit dynamics demonstrated that a network with strong excitatory-inhibitory balance, capable of CA3-driven reactivation, is consistent with our experimental results. Together, these results show that engram population dynamics are shaped by spontaneous states, behavior, and memory.

## Introduction

While decades of research have established the central role of the hippocampus in episodic memory formation, the precise mechanisms by which neuronal populations are selected to participate in individual memories are still undefined. A rich body of literature suggests that learning drives plasticity within populations of neurons activated by experience.^1,2^ This plasticity involves a dynamic series of changes in gene expression and synaptic connectivity which are thought to drive corresponding changes in activity patterns at the population level. Together these changes in gene expression, connectivity, and activity are thought to constitute a neurophysiological memory trace and are often referred to as a memory engram.^1^

Historically, two complementary approaches have been used to characterize such engrams. First, in vivo population recording of cells active during memory tasks and spatial navigation has been fundamental in understanding the coding properties of hippocampal neurons.^3^ Second, ‘engram-tagging’ studies have used immediate early gene (IEG) expression, such as *Fos* or *Arc*, to identify putative engram cells and to demonstrate their necessity and sufficiency in driving the behavioral expression of memory.^4–6^ While numerous cases of artificial manipulation of engram populations have driven a suite of behaviors ranging from fear recall (i.e., freezing^5,6^), approach^7,8^ and motivated behaviors,^9,10^ surprisingly less is known about how patterns of activity in engram populations change with learning.

Bridging the gap between cellular mechanisms and behavioral memory expression requires linking fast, intracellular events such as gene expression and spontaneous activity changes to slower, systems-level outcomes like post-learning population dynamics and behavioral phenotypes. Although a few studies have recorded hippocampal engram-tagged cells during navigation after *Fos*-based tagging^11,12^—revealing distinct spatial selectivity and ensemble dynamics compared to non-engram cells—how the activity of engram populations transforms across the course of memory formation remains poorly understood.

Two complementary hypotheses inform how engram populations relate to memory formation. The allocation hypothesis proposes that pre-existing cellular excitability influences the recruitment of cells during novel experiences.^13,14^ This idea is supported by numerous experiments that manipulate cellular excitability levels *before* learning. In parallel, engram-tagging research has converged on the idea that coactivity among engram cells emerges *after* learning and therefore comprises a physiological correlate for memory.^15–17^ This hypothesis is supported by recent work identifying structural and synaptic properties of engram cells in vitro–namely, that engram cells show a higher level of connectivity between each other compared to non-engram cells.^17–20^

To evaluate these hypotheses, we used a combination of activity-dependent genetic tagging, longitudinal calcium imaging, and computational modeling to characterize population activity in CA1 engram populations before and after learning. First, we discovered that spontaneous activity during periods of rest before learning was informative for engram allocation up to two days before IEG expression. In contrast, engram cells were less modulated by velocity than non-engram cells, though both populations participated in sequential activity during head-fixed locomotion. These findings indicate that intrinsic dynamics and locomotion differentially influence engram dynamics.

Furthermore, we evaluated how correlated activity amongst engram cells changes with learning. We observed a subset of engram cells with increased pairwise correlations during rest, after contextual fear learning. Finally, we utilized trace conditioning and a conditioned stimulus-mediated (CS) recall paradigm to describe the effects of memory cues on CA1 engram dynamics. CS presentation resulted in increased population activity that was greater in engram cells across trials, in addition to a stable correlation structure between engram cells. These reactivation dynamics were consistent with network simulations incorporating strong excitatory-inhibitory balance and CA3 reactivation. Overall, our results bridge in vivo cellular dynamics with *Fos*-expressing populations, revealing that pre-existing biology, as well as state- and memory-dependent changes, govern the physiology of memory-engram cells across learning and memory.

## Results

### Spontaneous activity in engram cells before and after learning

To record engram dynamics over time, we used tetTag virus constructs to identify engram cells and transgenic Thy1-GCaMP6f mice to record calcium activity in dorsal CA1 (<u>Figure 1A-B</u>). During imaging, head-fixed mice were free to walk on a rotating disk and we measured spontaneous locomotion. We used a custom kinematic headplate^21^ to register the head across sessions and reliably record calcium activity (GCaMP) and tet-driven mCherry in the same populations across days (<u>Figure 1B</u>, <u>Supplementary Figure 2A</u>). In a 405 x 405 μm field of view we observed an average of 177 +/- 6 active neurons. We were able to track the same cells across days and observed a turnover in the pool of active cells registered across 4 days (<u>Figure 1F</u>). 55.0% +/- 1.7% of cells from Day 0 remained active on Day 4, while 45.0 % +/- 1.7% percent of cells turned over from Day 0 to Day 4. Tet-driven mCherry expression was used to identify putative engram cells (<u>Figure 1C</u>-D, <u>Supplementary Figure 1</u>, Methods).

**Figure 1.**
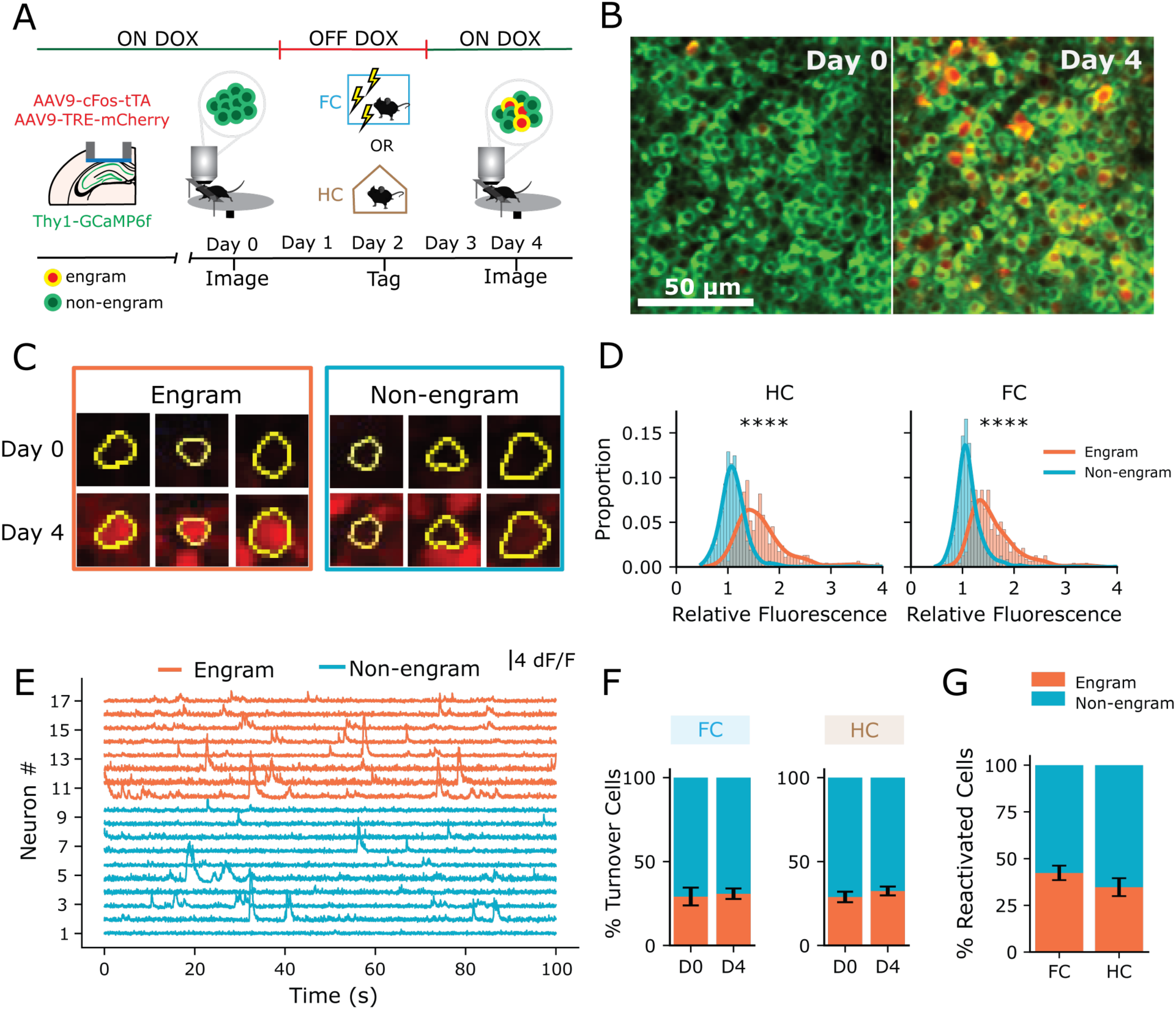
Imaging spontaneous activity in engram cells before and after learning. A) Experimental timeline. Doxycycline (Dox) is removed after imaging on Day 0 to open mCherry tagging window. Mice are off dox Days 1-2 and either fear conditioned on Day 2 or remain in homecage. Dox is replaced on Day 2 and spontaneous imaging of the same CA1 population is repeated on Day 4. B) Example dual color FOV before and after tagging. C) Representative engram and non-engram cells before and after mCherry tagging. D) Relative fluorescence of mCherry expression in engram vs non engram cells on Day 4 (engram vs non-engram cells: HC *p=*5e-115 n=1357 cells, FC *p=*2e-113 n=1214 cells) E) Representative calcium traces of spontaneous activity in engram and non-engram cells F) Percent of cells active on D0 only (OUT cells) and D4 only (IN cells). 2Way RM ANOVA Group x Session, Group: *p=*0.89, Session *p=*0.21, Interaction *p=*0.65. N=8 mice FC, N=8 mice HC. G) Percent of each cell type stably active on both days. FC 42.4% vs HC 34.7%, t-test *p=*0.25, N=8 mice FC, N=8 mice HC.

To assess how the activity of engram cells changes over time, we recorded CA1 populations before and after learning. Day 0 baseline spontaneous activity was recorded in a population of CA1 pyramidal neurons prior to *Fos*-tagging (<u>Figure 1A</u>). On Day 2, mice either underwent a 3-shock contextual fear conditioning protocol (FC, n=8, <u>Supplementary Figure 3C</u>) or remained in the homecage (HC, n=8), a neutral experience that still drove *Fos-*tagging. Dox was replaced on Day 2 to close the window for mCherry labeling, and a second imaging session of spontaneous activity took place on Day 4 (<u>Figure 1A-B</u>).

We classified cells as either engram or non-engram based on their Day 4 mCherry expression (<u>Figure 1C-D</u>, see Methods). Therefore, we use the terminology ‘engram’ here, to categorize cells in both FC and HC groups identified by their mCherry-expression in a given tagging window (<u>Figure 1A-C</u>). Approximately ∼35-40% of our recorded cells on both days were mCherry+ and therefore classified as engram cells (<u>Figure 1E</u>, <u>1G</u>). This proportion was similar between in vivo imaging and histological sections (<u>Supplementary Figure 3A-B</u>). In addition to cells active on both days, we defined two other classes of cells: those only active on Day 0 and cells only active on Day 4, which were comparable across groups (<u>Figure 1F).</u>

### Increased activity prior to tagging biases Fos-tagging

We first sought to test a prediction of the allocation hypothesis, namely if spontaneous activity before learning could predict engram tagging. To do this, we deconvolved calcium activity and calculated event rates for each cell on Day 0 (Methods). Within each animal, cells in different activity deciles were distributed in space throughout the FOV (<u>Figure 2A-C).</u> We observed a significant correlation between a cell’s activity decile on Day 0 and the likelihood it would be *Fos*-tagged (<u>Figure 2D</u>, *R*^2^=0.82, *p=*0.00033). Interestingly, this effect was seen in both FC and HC groups (<u>Figure 2E-F</u>, FC *R*^2^*=* 0.65, *p=*0.0048, HC *R*^2^ *=* 0.77, *p=*0.0009). These results suggest that increased activity up to 48 hours before *Fos*-tagging correlates with allocation, independent of which experience was tagged (i.e. aversive footshock or neutral homecage). Notably, spontaneous activity during rest, but not during locomotion, was correlated with engram tagging (<u>Figure2G-I</u>, Rest: *R*^2^ *=* 0.8, *p=*0.00049; Run: *R*^2^ *=* 0.31, *p=*0.09). However, in HC mice, the run activity was still weakly correlated with *Fos*-tagging (<u>Figure 2I</u>, *p=*0.01).

**Figure 2.**
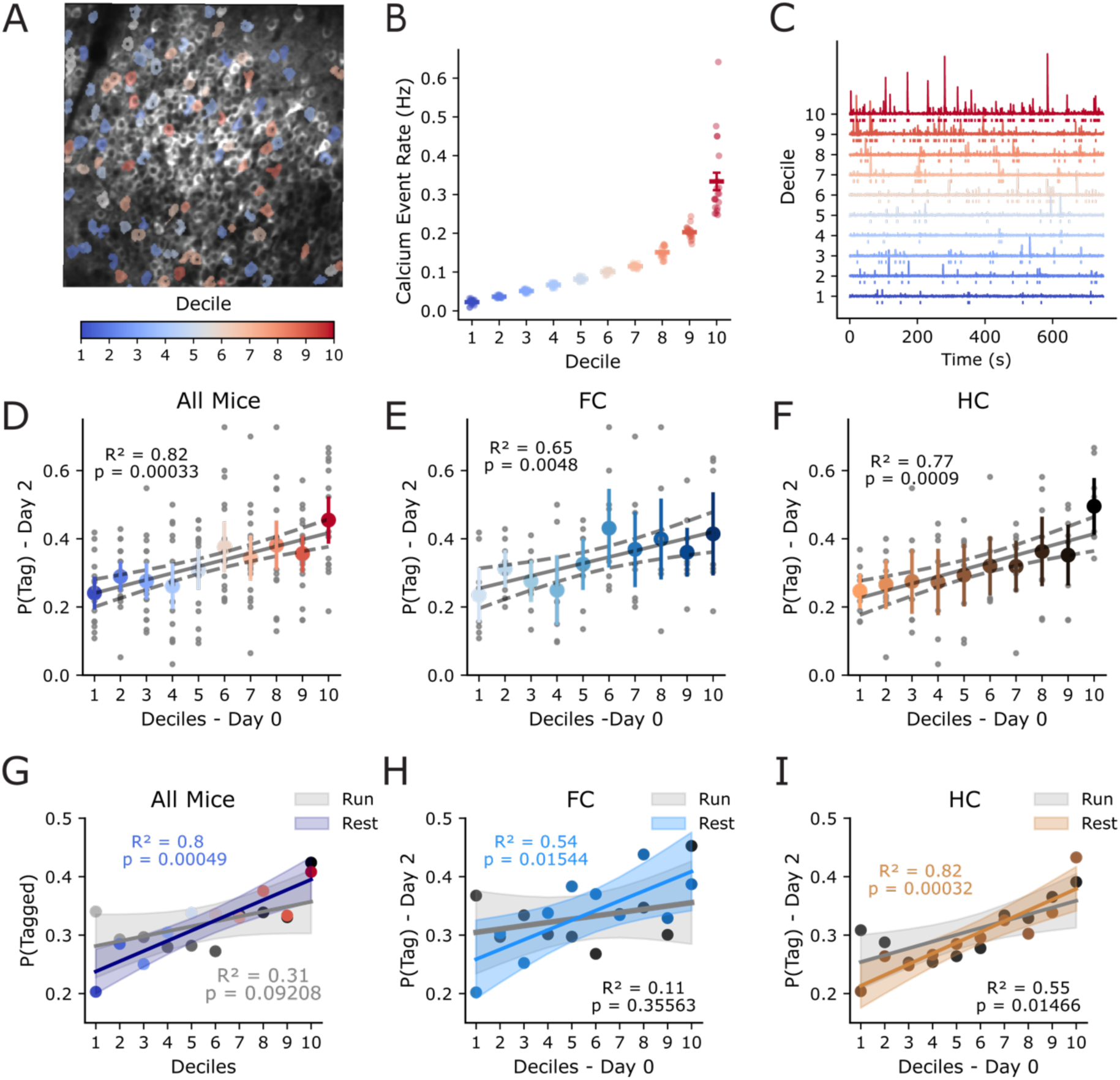
Day 0 spontaneous activity predicts engram allocation two days before tagging. A) Distribution of cells in each decile throughout the example FOV. ROIs are colored by decile (blue = lower activity, red = higher activity). B) Example animal with cells split into deciles. C) Example dF/F traces and calcium events from example cell in each decile. D) Linear regression of baseline calcium activity deciles and probability of Fos-tagging. Blue colors: least active, red: most active. All mice (FC N=8 and HCN=8), N=16, *R*^2^ = 0.82, *p=*0.00033. E) FC mice regression of baseline calcium activity deciles and probability of *Fos*-tagging. light colors: least active, dark colors: most active. N=8; *R*^2^*=*0.64, *p=*0.0048. F) HC mice regression of baseline calcium activity deciles and probability of *Fos*-tagging. light colors: least active, dark colors: most active. N=8; *R*^2^*=*0.77, *p=*0.0009. G) Linear regression of baseline calcium activity split by behavioral state: Rest period activity (navy) or run period activity (gray) Rest: R^2^=0.8, *p=*0.00049; Run: R^2^=0.31, p=0.09 H) Same as G) but for FC mice only (N=7). Rest (blue) : R^2^ = 0.54, p=0.015; Run (gray): R^2^ = 0.11, p=0.35 I) Same as G) but for HC mice only (N=6). Rest (brown) R^2^ = 0.82, p=0.00032; Run (gray) R^2^ = 0.55, p=0.015.

To validate these results, and since this analysis averaged information across cells and animals, we also used hierarchical Bayesian generalized linear models to test the relationship between Day 0 activity and likelihood of engram tagging (HB-GLM; <u>Supplementary Figure 5A</u>, <u>Methods</u>). This analysis revealed a strong relationship between Day 0 activity during rest and tagging, consistent with the previous analysis (<u>Supplementary Figure 4C</u>). More specifically, GLMs that included resting state activity rates demonstrated improved predictive performance against a model only based on running state activity rates, or a null model (<u>Supplementary Figure 4B-D</u>).

Together, these analyses of Day 0 activity rates provide converging evidence that spontaneous activity in engram cells during rest is predictive of future engram recruitment. This supports the idea that internal, pre-experience dynamics, particularly evident during rest, are a critical signature of engram allocation.

### Engram cells remain more active during rest across conditions

We next asked if engram cells maintained their higher activity across days, and if this changed after fear learning. To address this, we registered cells across both Day 0 and Day 4 (<u>Supplementary Figure 2</u>) and compared their mean calcium event rate across each session. Interestingly, engram cells had higher activity rates at both timepoints in both FC and HC mice (<u>Figure 3A</u>), in agreement with previous recordings of engram cell activity in CA1.^11,12^ We saw no increase in activity amongst engram cells from Day 0 to Day 4 in either group of mice, but found a Group x Session interaction (<u>Supplementary Figure 5A</u>, *p=*0.001 Linear Mixed Effects Model). To control for variability in locomotion (<u>Supplementary Figure 5B)</u> and the general increase in rates caused by more locomotion (<u>Supplementary Figure 5C</u>), we recalculated event rates during separate periods of locomotion and periods of quiet rest (<u>Figure 3B-C</u>). We observed a modest decrease in event rate across both groups during resting activity on Day 4 (*p=*0.03) and not running (*p=*0.85, <u>Figure 3B-C</u>). Interestingly, the increased activity in engram cells was only significant during rest periods (<u>Figure 3B</u>, *p=*0.004) and not run periods (<u>Figure 3C</u>, *p=*0.145). Overall, these results suggest that *Fos*-identified engram populations display higher levels of spontaneous activity that was more accurately detected during periods of quiet rest compared to running.

**Figure 3.**
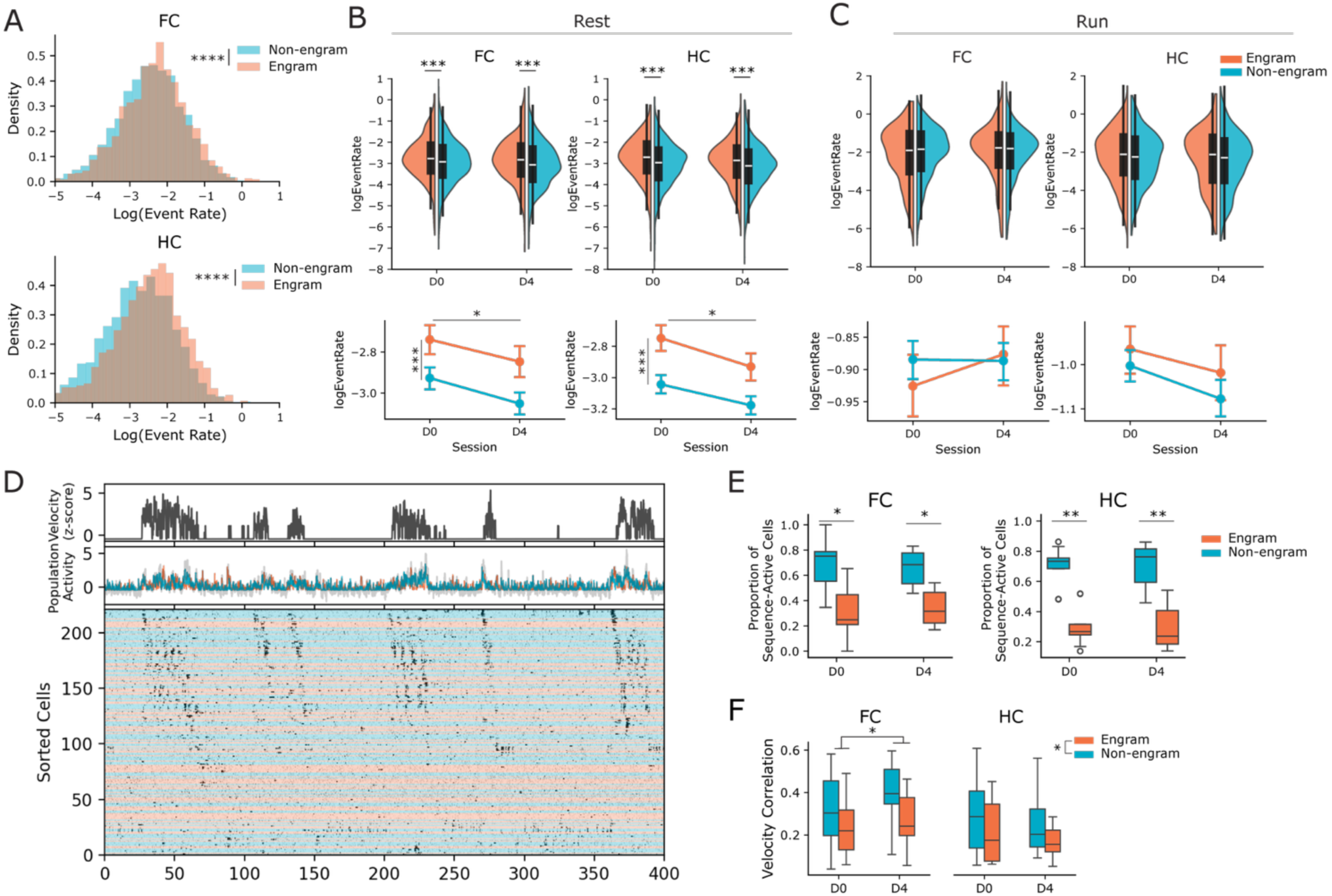
Engram and non-engram populations are differentially modulated by behavior state. A) Distribution of calcium event rates for engram and non-engram cells for each group. Linear mixed effects model, Population x Session. Main effect of Population, *p=3e-9* Session *p=*0.18. FC: N=8 mice, n=2384 cells, HC N=8 mice, n=2379 cells. B) Average calcium event rates during rest periods across sessions for each population. Linear mixed effects model, Effect of Population p=0.004, Session p=0.026, Group p= 0.198. FC: N=7 mice, n=2183 cells, HC N=6 mice, n=1894 cells. C) Average calcium event rates during run periods across sessions for each population. Linear mixed effects model, no significant main effects or interactions. Main effect of Population p=0.145, Session p=0.875, Group p=0.430. FC: N=7 mice, n=2183 cells, HC N=6 mice, n=1894 cells. D) Example rastermap sorting of sequential activity associated with locomotion bouts. Top: normalized velocity. Middle: Summed population activity for engram and non-engram populations. Bottom: sorted rasterplot from rastermap, color-coded by cell type. E) Proportion of sequence-participating cells from each population in FC and HC groups. 2Way RM ANOVA, Population x Day. FC Population *p=*0.0096, Interaction *p=*0.38, HC Population *p=*0.0005, Interaction *p=*0.91. Pairwise t-tests, FC (N=12 FOVs), HC (N=10 FOVs). F) Velocity correlation of each population. Linear Mixed Effects Model Group x Day x Population. Main effect of Population p=0.047, Group x Day interaction p=0.028. FC (N=12 FOVs), HC (N=10 FOVs).

### Engram and non-engram cells are differentially modulated by behavior state

Previous studies have identified spatially or temporally patterned sequences of hippocampal activity that are associated with navigation, memory and locomotion.^22–24^ These sequential dynamics are thought to organize memories by providing biological scaffolds for space and time that are relevant to behavior. We asked how engram and non-engram cells may contribute to such dynamics. Examination of population dynamics from CA1 with Rastermap^25^ revealed sequential patterns of activity that were consistently associated with locomotion bouts (<u>Figure 3D</u>). While both engram and non-engram cells participated in these sequences, a larger proportion of sequentially active cells were non-engram cells (<u>Figure 3E</u>, 2-Way RM-ANOVAs Population x Day, Effect of Population, FC: *p=*0.0096 HC: *p=*0.0005). Furthermore, while fear conditioning resulted in an increased correlation between activity of both cell types and velocity (Linear Mixed Effects Model, *p=*0.028), the overall population activity of non-engram cells was more correlated with velocity across conditions (<u>Figure 3F</u>, Linear Mixed Effects Model, *p=*0.047). These data suggest that while engram and non-engram populations participate in locomotion related sequences, engram populations are less modulated by locomotion, which is consistent with their having higher activity during rest states.

### Subsets of engram cells increase their spontaneous correlations after fear learning

A key prediction of many engram-tagging studies is that after learning, engram neurons have stronger synaptic connectivity that drives their co-activity.^1,26^ To test this prediction, we calculated pairwise Spearman correlations for all pairs of cells on Day 0 and Day 4 (<u>Figure 4A),</u> such that each pairwise correlation was between engram cells (E/E pairs), engram and non-engram cells (E/N) or non-engram cells (N/N). After fear learning, mean pairwise correlations did not differ significantly across populations, although FC mice showed a nonsignificant trend toward lower E/E correlations (<u>Supplementary Figure 6D</u>, Linear Mixed Effects Model Group x Population, *p=*0.07). After separating out locomotion state, this effect was driven by changes in running behavior across days in FC mice (<u>Supplementary Figure 5C)</u>, and disappeared when calculating rest and run correlations while controlling for time spent in each state (<u>Supplementary Figure 6E</u>).

**Figure 4.**
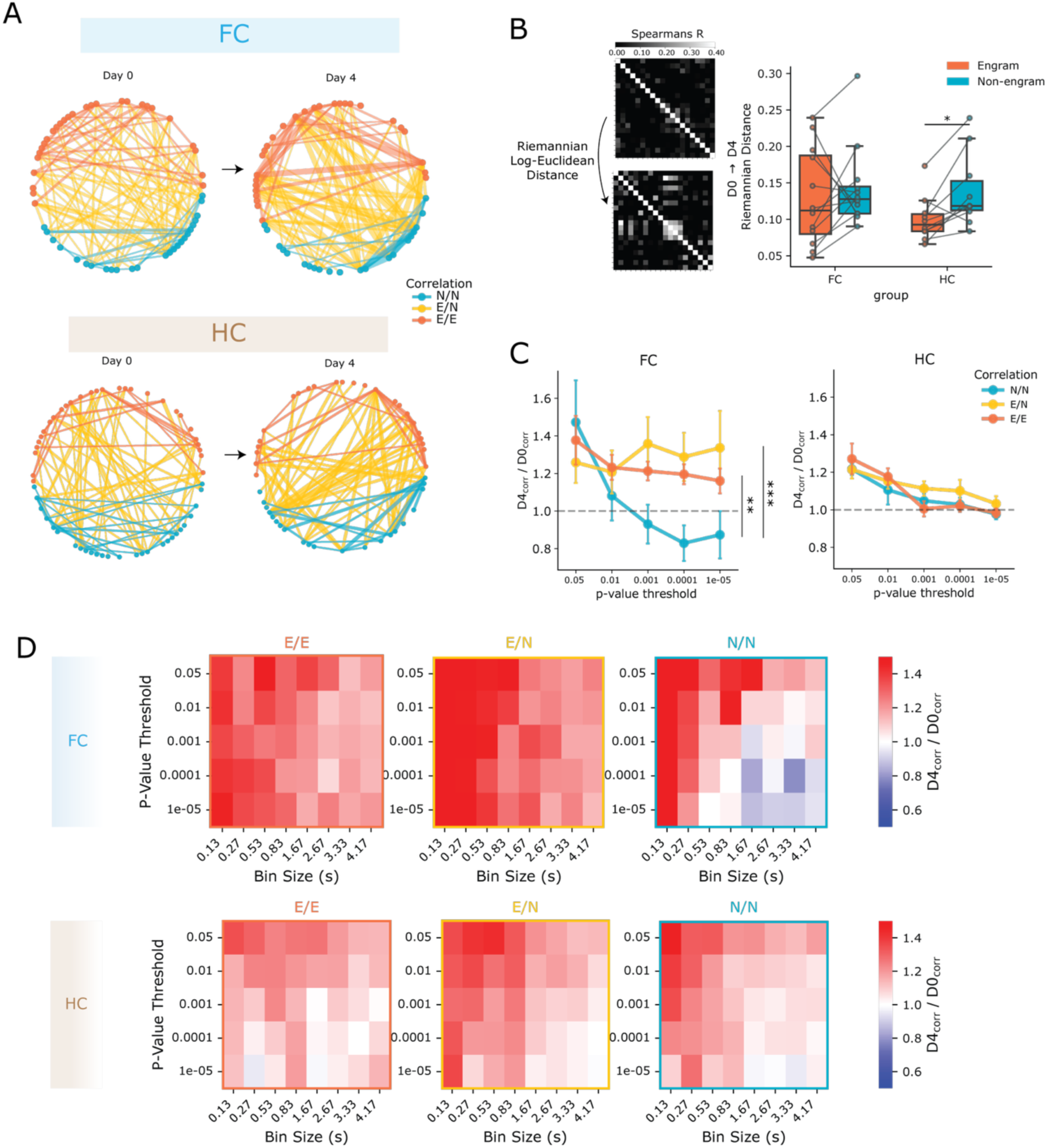
Subsets of engram correlations increase during rest after fear learning. A) Example network graphs displaying Spearman correlations between cells from one FC and one HC mouse. Left: Day 0, Right: Day 4. For simplicity, the top 2% of correlations are shown as edges between nodes, scaled by their correlation strength. Orange nodes indicate engram cells, teal nodes indicate non-engram cells. Orange edges = engram/engram (E/E) correlations, yellow edges = engram/non-engram (E/N) correlations, teal edges = non-engram/non-engram (N/N) correlations. Cell ID sorting is consistent across both days. B) Riemannian log-Euclidean distance of E/E and N/N correlation matrices from D0 to D4. Linear Mixed Effects Model Effect of Population p=0.432, Group p=0.198, Interaction p=0.310. Post hoc Wilcoxon signed-rank tests, HC *p=*0.039, FC *p=*0.57. FC N=12 FOVs HC N=10 FOVs. C) Normalized change in correlations for thresholded cell pairs across significance levels. A bin size of 1.7 seconds was used. Linear Mixed Effects Model, Group x Population interaction. E/E x FC and E/N x FC interactions *p=*0.002 and *p=*0.0005 respectively. N=7 mice HC, N=6 mice. D) Heatmap of normalized change in correlation for thresholded pairs across timescales (x-axis, bin sizes) and significance thresholds (y-axis, p-values). Red: increase in correlation across days, Blue: decrease across days, white: no change. N=7 mice HC, N=6 mice.

We explored the potential mechanisms underlying this result after learning by implementing a spiking network model of CA3-CA1 (see <u>Methods</u>), based on a Hebbian increase in synaptic weights between engram-units in both regions (<u>Supplementary Figure 9A</u>). Initially, increasing synaptic weights in our model led to increases in pairwise correlations between engram units (<u>Supplementary Figure 9B</u>). However, the addition of homeostatic plasticity and strong global excitatory-inhibitory balance throughout the network, was sufficient to mask this mean change in engram correlations, similar to our in vivo data (<u>Supplementary Figure 9C-D</u>). This points to E/I balance as a potential circuit mechanism for maintaining balanced coactivity across populations that is consistent with our observed spontaneous activity here.

To see how network as a whole changed with learning, we analyzed geometric distance of the correlation matrices from Day 0 to Day 4 using a Riemannian (log-Euclidean) distance metric.^27,28^ Since correlation matrices are positive semi-definite, this geodesic metric is better than Euclidean distance at capturing similarity of the populations’ coactivity. In HC control mice, the engram matrices had reduced distance across days compared to the non-engram matrices (<u>Figure 4B</u>, paired t-test *p=*0.03). In contrast, with fear learning, FC engram and non-engram correlation matrices had comparable distances (<u>Figure 4B</u> paired t-test *p=*0.96). This indicates that at baseline, *Fos*-expressing engram networks demonstrate more stable co-activity structure across days compared to non-engram *Fos*-cells, but that fear learning drives reorganization of both engram and non-engram co-activity structure.

We next asked how subsets of co-active cells contributed to this change. To do this we analyzed only significantly correlated cell pairs by applying significance thresholds to our pairwise correlations (<u>Supplementary Figure 6A-C).</u> Correlated cell-pairs were thresholded such that only pairs with significant correlations on both days were analyzed (p-values less than thresholds ranging from 0.05 to 10e-5, <u>Supplementary Figure 6A-C</u>). In addition, to account for time lags between pairs of cells, we repeated these correlations on binned traces of the deconvolved activity using a range of time bins (0.27 to 4 seconds). We found that during rest, in FC mice, E/E and E/N cell pairs became more correlated after fear conditioning compared to N/N pairs, <u>(Figure 4C-D</u>). This effect was not observed in HC mice (<u>Figure 4C-D</u>, Linear Mixed Effects Model, Group x Population interaction, FC x E/E *p=*0.002, FC x E/N *p =*0.0005). Conversely, correlations during running did not show these effects (<u>Supplementary Figure 6F-G</u>).

In summary, these data suggest that a subset of existing engram correlations increase after learning in a state dependent manner, and co-activity structure between engram cells is reorganized. Consistent with our activity rate results, these changes in engram cell correlations are revealed during states of quiet rest.

### CS presentation drives increased activity and stable correlation structure in engram cells

While our data thus far demonstrate how spontaneous activity dynamics evolve across memory formation, we next sought to explore how memory retrieval states could impact activity in engram populations. To do this, we tracked engram population activity in a new group of mice that underwent trace fear conditioning followed by imaging engram population activity during the presentation of the tone conditioned stimulus (i.e., CS+) (<u>Figure 5A</u>, <u>Supplementary Figure 7</u>). Similar to our first experiment, baseline activity was measured on Day 0 and dox was removed. While off dox, trace fear conditioning (TFC) training consisted of 3 trials of a 2.8 kHz tone (20s), followed by a 20s trace period, and an aversive foot shock. In a separate cohort of mice, we ensured that this tone CS served as an effective fear cue to induce a fear response in a distinct, neutral environment (<u>Supplementary Figure 3E</u>). After training, mice were again imaged on Day 4 for a recall session. After 10 minutes of baseline (pre-CS+) recording, 10 trials of the CS+ were presented, once every 100 seconds for the remainder of the session (<u>Figure 5A</u>). This approach enabled us to record CA1 engram cells during a baseline period after learning, as well as during CS+-evoked activity.

**Figure 5.**
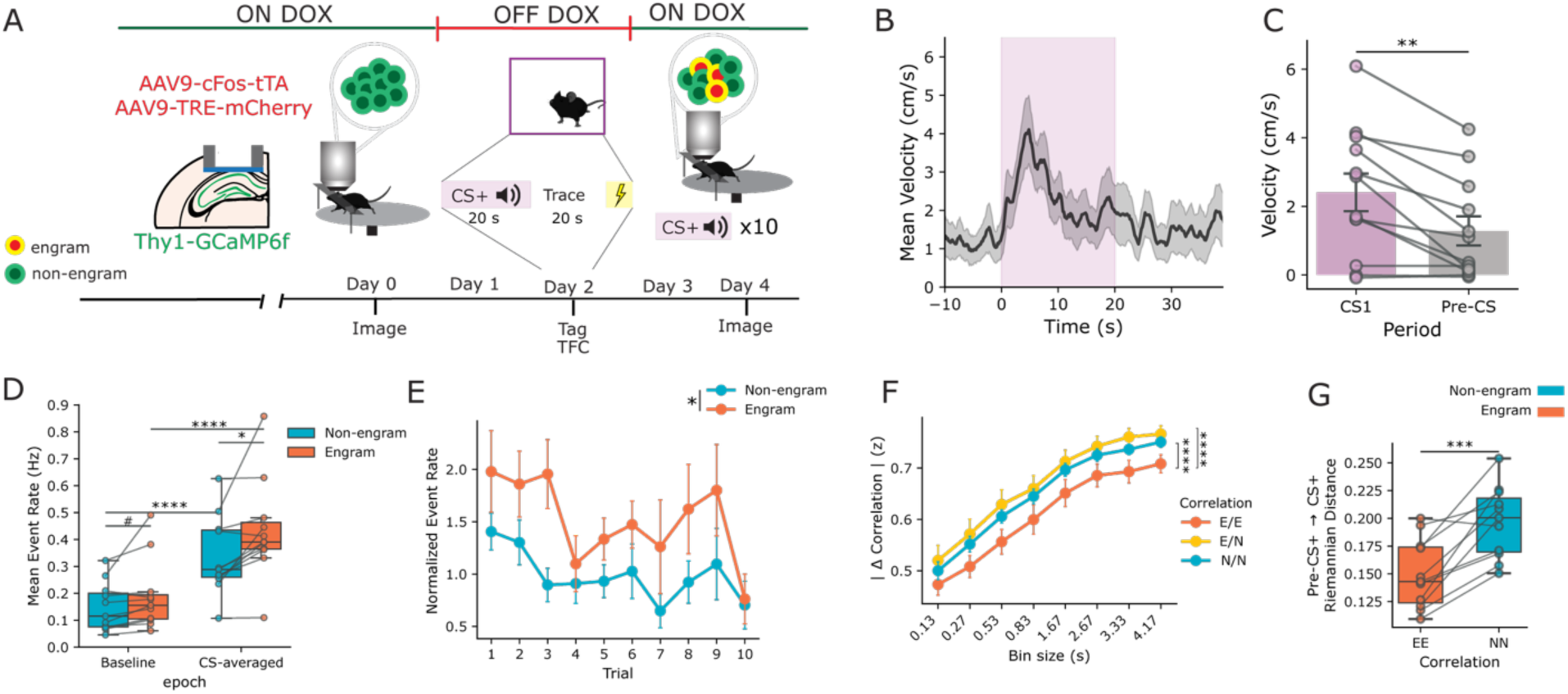
Engram activity during cued reactivation. A) Schematic of experiment timeline for recording engram CS reactivation after trace fear conditioning. B) Trial averaged velocity trace upon CS presentation during imaging on Day 4, Recall. N = 11. C) Animal-wise increase in average velocity during CS presentation. Paired t-test *p=*0.005, N=11. D) Increase in event rate among engram cells persists during CS presentations. 2way RM ANOVA Population x Epoch, Population *p=*0.002, Epoch *p=*1e-6, Population x Epoch *p=*0.03. Engram vs Non-engram paired t-tests: Baseline *p=*0.06, CS-averaged *p=*0.03. Baseline vs CS+-averaged paired t-test: Non-engram *p=*4e-6, Engram *p=*4e-6. N=11 mice. E) Event rates across trials in engram and non-engram cells, normalized to pre-CS+ period. 2Way RM ANOVA Population x Trial. Population *p=*0.01, Trial *p=*0.03, Population x Trial: *p=*0.18, N=11 mice. F) Normalized absolute value of change in pairwise correlation from pre-CS+ to CS+/Trace periods. Linear mixed effects model, Population + Bin size, Effect of E/N, *p=*1e-11; N/N, *p=*3e-6. N=11 mice. G) Riemannian distance between E/E and N/N correlation matrices from pre-CS+ to CS+/Trace periods. Paired t-test, *p=*0.0005. N=11 mice.

First, we replicated our result from our spontaneous recordings (<u>Figure 2A-B</u>)—we found that during the baseline period, engram cells demonstrated higher activity rates during rest periods (<u>Supplementary Figure 8B-C</u>). In addition, we still found locomotion-related sequences during CS trials, similar to before CS presentation (<u>Supplementary Figure 7A-B</u>). Upon CS+ presentations, we observed increased locomotion in head-fixed mice, which often began a new running bout from rest (<u>Figure 5B-C</u>, paired t-test *p=*0.006) consistent with a learned US-CS association (but see Discussion). CS+ presentation drove increased activity in both engram and non-engram cells (2Way RM ANOVA Main Effect of Epoch, *p=*1e-6, Population x Epoch 0.027) with a larger increase in engram cells (<u>Figure 5D</u>, paired t-test, *p=*0.028). While activity rates varied across trials, engram cells consistently demonstrated higher activity than non-engram cells (<u>Figure 5E</u>, 2Way RM ANOVA, Effect of Population *p=*0.01). Finally, our spiking network model was also consistent with these reactivation data, when we simulated upstream engram reactivation in CA3 (<u>Supplementary Figure 9D</u>). In this simulation, when a subset of CA3 engram units received excitatory input, mimicking the recall stimulus (<u>Supplementary Figure 9E, 9G).</u> As a result, downstream CA1 engram units increased their spiking relative to non-engram units (<u>Supplementary Figure 9F-G</u>), similar to our in vivo data (<u>Figure 5D-E</u>). Our network model recapitulated these results when targeted stimulation of excitatory engram cells was paired with a nonspecific disinhibitory input <u>(Supplementary Figure 9E</u>).

To account for stimulus-specific effects, a separate cohort of mice (n=5) underwent an additional day of imaging (<u>Supplementary Figure 8A),</u> when they were presented with interleaved trials of the fear CS (CS+) and a neutral auditory CS (CS-, pulsed 12 kHz tone). Again, before any CS presentation engram cells had higher activity rates, and running increased activity of both engram and non-engram populations similarly (<u>Supplementary Figure 8D</u>, Linear Mixed Effects Model Effect of Run *p<*0.0001, Population p=0.03). However, in the trace period following CS+ trials, engram cell activity during locomotion was significantly higher than non-engram cells, an effect that was specific for the CS+ (<u>Supplementary Figure 8E</u>, Linear Mixed Effects Model, Population x CS x Run *p=*0.016). This was not simply driven by higher running velocity during CS+ versus CS-trace periods (<u>Supplementary Figure 8F-G</u>).

Finally, to see how CS presentation affected correlations between engram cells, we repeated our correlation analyses, comparing pairwise correlations during the CS+ and trace periods to baseline, before any CS. We took cells with significant Spearman correlations (*p<*0.05), and calculated the change in their correlation coefficient from the baseline (pre-tones) to the CS+/trace periods. Surprisingly, when we compared the absolute value of this change in correlation across populations, the magnitude of this change for E/E pairs was lower than E/N and N/N pairs (<u>Figure 5F</u> Linear Mixed Effects Model, Main Effect of N/N and E/N, *p<*0.0001). These results suggest that upon CS presentation, E/N and N/N pairs demonstrate stronger reorganization of their co-activity patterns than E/E. This was further supported by Riemannian distances of E/E versus N/N correlation matrices from the baseline period to CS/trace periods. E/E co-activity demonstrated a reduced Riemannian distance (<u>Figure 5G</u>, paired t-test, *p=*0.0005), suggesting engram cells maintained a stable co-activity pattern upon CS+ presentations, while N/N co-activity reorganized more.

In sum, our results show that the CS+ causes engram cells to increase activity relative to non-engram cells, while maintaining a stable correlation structure with each other. In contrast, non-engram cells have lower activity rates and larger changes in their correlations, both with each other and engram cells.

## Discussion

We investigated the dynamics of *Fos*-tagged engram cells in CA1 before and after fear learning using an intersectional approach that combined in vivo imaging of cell populations, activity dependent tagging and spiking models. Overall, our experiments reveal that engram calcium activity is modulated by pre-existing hippocampal dynamics, behavioral states, and fear learning.

First, we observed that activity up to two days before fear conditioning was predictive of *Fos*-tagging (<u>Figure 2</u>). This spontaneous activity was most informative about cell identity when including periods of quiet rest, a state in which engram cells displayed increased calcium activity in spontaneous recordings (<u>Figure 2</u>, <u>Figure 3A-C</u>). We also found that sequential hippocampal activity during locomotion recruited both engram and non-engram cells, however, engram cells were less strongly modulated by velocity (<u>Figure 3D-F</u>). Next, we discovered that pairwise correlations during rest states increased in a subset of engram neurons after fear learning, an effect that was learning-specific (<u>Figure 4C-D</u>). Finally, recording during CS+ cued reactivation demonstrated increased activity (<u>Figure 5D-E</u>) and stable correlation structure (<u>Figure 5F-G</u>) amongst engram cells during memory reactivation. Together, these results reveal several novel aspects of the underlying activity of *Fos*-tagged engram populations as it relates to behavior and memory.

### Allocation revealed by spontaneous dynamics

Our finding that increased spontaneous activity prior to experience is predictive of IEG-tagging further supports the popular allocation model of engram formation.^13,14^ Previous evidence for the allocation model came initially from manipulation studies, in which over-expression of CREB or optogenetic stimulation of cells prior to learning increased their likelihood of recruitment and reactivation during memory recall.^29–33^ While there is broad support for this model of engram formation, several aspects of the allocation model have yet to be characterized.

First, while changes in excitability are proposed to be the key factor enabling allocation, there is no clear consensus on the endogenous timescales over which intrinsic excitability fluctuates in the hippocampus. On the one hand shorter timescales on the order of minutes to hours have been observed to contain physiological activity predictive of memory allocation^30,33^, whereas other studies propose that intrinsic fluctuations that can exist across days, as a mechanism for linkage of fear memories across days.^34–38^ The timescale of our recordings is consistent with the latter view, suggesting that activity of some cells may fluctuate on days-long timescales.

Second, it remains unclear whether allocation to an engram population is cell autonomous, such that each cell’s probability of recruitment is independent of others, or whether pre-existing networks are recruited together.^39^ Several lines of evidence offer support for the latter. One study implemented longitudinal spine imaging to show that CA1 neurons with less spine turnover in the days leading up to an environmental enrichment experience were recruited and expressed *Arc*, which the authors interpreted as a reflection of an underlying stable ensemble that is strengthened by experience.^19^ Consistent with this, in vitro measures of hippocampal *Fos*-tagged engram cells showed that their co-activity was not completely abolished by post-experience anisomycin^17^, consistent with a degree of pre-existing connectivity between engram cells. Finally, a more recent study found that excitability of place cells was predictive of their stability across days^40^, which parallels what we observe here: just as high-activity place cells maintain stable fields over days, we observe a strengthening of co-activity between engram cells with elevated pre-learning activity. Together, we suggest that elevated activity observed during rest days prior to experience may therefore be another signature of such pre-existing networks, which have been observed in other conditions of hippocampal activity (i.e. preplay).^41^

To address these open questions, future research into engram ensembles should focus on characterizing endogenous fluctuations in the intrinsic physiology of networks at long timescales *prior* to learning. A recent study used in vivo imaging to track endogenous fluctuations in calcium dynamics prior to IEG-tagging,^33^ finding that cells with higher activity in the minutes-hours before fear conditioning were more likely to undergo IEG-tagging, which resonates with our findings. However, this study reported that increased excitability was only detectable in engram cells within the same day as learning, which differs from our result. Notably, the analysis in this study normalized calcium activity of individual cells across days, while here we compared the engram- to non-engram population activity, which does not preclude additional changes *within* the engram population on the day of learning. Taken together, evidence across multiple groups points to the existence of heterogeneous dynamics, including both short and long timescale intrinsic fluctuations. Such heterogeneity could be rooted in various biological underpinnings including, but not limited to, intrinsic CREB expression, receptor recycling and homeostatic structural-functional changes.

### Fos ensembles for aversive and neutral experiences

In our analyses of baseline activity, we found the relationship between prior spontaneous activity and engram-tagging existed in both aversive (FC) or neutral (HC) conditions (<u>Figure 2</u>, Day 0). This suggests that increased spontaneous activity predicts *Fos*-tagging, regardless of experience valence. Interestingly, however, the correlation between activity decile and likelihood of tagging was stronger in the HC mice (<u>Figure 2E-F</u>, R^2^=0.82 vs R^2^=0.54). This suggests that in FC mice, other factors aside from intrinsic activity levels (e.g. dynamics during the learning itself) determine allocation to the *Fos* population when a fear memory is formed. HC mice still had a weaker relationship of activity decile with tagging during running states, while FC mice did not (<u>Figure 2H-I).</u> Together, these findings reveal that both behavioral state, and experience (FC vs HC) influence the correlation between endogenous activity level and recruitment to an engram ensemble.

In prior work studying their behavioral relevance, FC-tagged cells drove freezing while neutral-tagged cells did not.^5,6,42,43^ While this approach focused on the role of valenced experiences in mediating the ability of *Fos*-tagged ensembles to drive behavior, the physiological similarities or lack thereof between neutral or fear-related *Fos* assemblies remained unmeasured. Our findings highlight two important considerations: first the importance of continued investigation into how learning-related changes in IEG-populations differ from the baseline physiology of *Fos*-ensembles involved in neutral or homecage experiences; and second, the need for experience-based behavioral controls in engram-tagging experimental designs. Our results here suggest that some features of IEG-ensemble activity may decouple behavioral experience from physiology of engram cells by identifying aspects of *Fos*-ensemble function that are independent of experience or valence.

### State-dependent dynamics in engram populations

Interestingly, our finding that increased spontaneous activity was important for future IEG-tagging was state-dependent, and strongest during rest-period activity. This observation is particularly relevant given that the hippocampus displays highly structured dynamics such as sharp-wave ripples (SWRs), replay and preplay during rest and offline periods.^22,23^ Interestingly, there is preliminary evidence that *Fos*-tagged CA1 pyramidal cells are strongly modulated by SWRs.^44^ Such structured activity is thought to support successful encoding and consolidation of episodic experiences, and contribute to memory guided decision-making.^22,45^ Alternatively, the preferential emergence of engram-specific differences during rest could reflect that locomotion broadly elevates activity across the population, masking underlying differences in baseline activity levels. Distinguishing between these possibilities will require future approaches capable of resolving fast-timescale sequential dynamics. Indeed, one interesting future direction would be to investigate whether engram cell assemblies display reliable and specific fast-timescale sequences of firing prior to learning and tagging, and whether these sequences change after learning.

While differences in engram vs non-engram dynamics were most evident in spontaneous activity during rest periods, during run periods both engram and non-engram cells participated in behaviorally relevant sequences of activity (<u>Figure 3D</u>). These behavioral-timescale sequences of activity are similar to previous reports that found repeated sequences of distance-coding in head-fixed mice with no visual cues.^46^ However, non-engram cells were more strongly correlated with velocity (<u>Figure 3E-F</u>). In addition, during spatial navigation in virtual reality, *Fos*-positive place cells also participated in sequential place field activations.^12^ These findings suggest that *Fos*-ensembles can participate in both spontaneous activations during rest and locomotion during navigation, which supports their involvement in both types of hippocampal activity.

### Subsets of engram cells strengthen their correlations after learning

Our experiments also revealed specific post-learning changes in functional correlations between CA1 engram cells. While we did not detect widespread increases in spontaneous correlated engram activity after learning in head-fixed mice, we did discover a subset of highly-correlated engram cells that strengthened their co-activity (<u>Figure 4A</u>, 4C-D). Our results are in agreement with a previous study in which a decrease in average correlation after learning was observed, in addition to the emergence of a subset of highly correlated cells after fear learning.^47^ Interestingly, the effect we observed extended to engram/non-engram correlations (<u>Figure 4C-D</u>), which is consistent with a recent connectomics study that also found increased synaptic strength both within engram/engram and engram/non-engram connectivity from CA3 to CA1.^20^ Finally, these increases were state-dependent as this effect was not seen in run-state activity.

These findings point to several novel insights: first, subsets of highly correlated engram cells demonstrate state- and learning-dependent changes across a days-long timescale in spontaneous activity. Second, the consequences of plasticity amongst hippocampal engram populations on circuit dynamics cannot be solely explained by Hebbian potentiation, since these correlations were state-dependent and also strengthened between engram and non-engram cells. This conclusion is supported by our model simulations, in which purely Hebbian strengthening of engram unit connectivity had to be constrained by increased global inhibition to align with in vivo results. Finally, the fact that this increase was only evident in cells with some existing degree of coupling is consistent with the idea that pre-existing functional relationships may shape memory ensembles and be stabilized during learning, and not necessarily emerge de novo. Future studies and analytical approaches may be aimed to identify these ‘sub-ensembles’ within IEG-labeled populations,^16,33^ and characterize how their existence or stability are changed by learning.

### Engram activity during CS+ presentation

Lastly, aside from spontaneous activity, we investigated how reactivation of the fear representation influenced population activity. We found engram cells were reactivated during CS+ presentation after trace fear conditioning, as CS+ presentations drove higher activity in engram cells compared to non-engram cells (<u>Figure 5D-E</u>). Importantly this difference in activity was not observed during neutral CS-trials (<u>Supplementary Figure 8E</u>). These results point to a competitive mechanism for memory retrieval in these populations, which has similarly been proposed as an aspect of memory allocation at the time of encoding.^30^ Our findings are the first to show that CA1 engram populations can be reactivated by the presentation of a fear-CS, reinforcing their participation in memory retrieval, consistent with *Fos*-reactivation studies.^15,48^

Beyond differences in event rate, pairwise correlations across the recall session revealed a surprising result: E/E pairs had smaller magnitude changes in their correlation strength upon CS+ presentation, in contrast with E/N and N/N pairs. This suggests that after learning, there is an existing level of coactivity between engram cells that remains stable during cued retrieval with a CS+. In contrast, during CS+, E/N and N/N correlations had stronger magnitude changes, suggesting their co-activity structure changes more drastically during CS+ presentation. These data represent a novel, in vivo, demonstration of engram versus non-engram coactivity during cued memory recall, and are consistent with a recent opto-tagging study.^49^ When stimulating CA1 *Arc*-tagged cells, this study reported a maintenance of endogenous cofiring patterns, similar to our findings during natural recall. While our CS+ protocol took place in a head-fixed preparation, it is possible that recording in freely moving mice where recall and encoding contexts are more similar, could yield distinct changes in co-activity.^50^ Future studies may use this approach to characterize different aspects of memory cue encoding, during learning or retrieval in *Fos* networks—such as higher-order analyses of engram population activity and/or remapping across extinction or new learning.

Another somewhat surprising result is that we did not observe sequential dynamics at stimulus onset or offset tiling the trace period, though the velocity modulated sequences still existed (<u>Supplementary Figure 7</u>). These results are consistent with a recent report that did not detect sequential activity in CA1 during head-fixed trace fear conditioning.^51^ This study did however find that subsets of neurons still encoded CS identity at longer timescales, which is similar to our finding that engram cells had distinct activity levels for CS+ versus CS-trials. These findings contrast with other learning tasks in which ‘time cells’ fired in reliable sequences in a task-dependent manner.^52–54^ These tasks generally incorporate shorter stimulus or delay durations, compared to trace fear conditioning, in addition to using appetitive learning. This discrepancy may indicate that longer-timescale delays in aversive conditioning may recruit distinct neural mechanisms,^55,56^ though additional experiments are warranted to explicitly investigate this.

### Limitations of the study

A key issue with many IEG-tagging studies lies in the difficulty of estimating the onset and duration of the tagging window. This problem is particularly acute with the tet-Tag system when doxycycline is administered in the animal’s diet. We chose this system for its wide use in engram-tagging studies and the evidence that cells labeled with this system can drive memory behaviors. However, several unknowns limit the precise knowledge of onset and duration of tagging including the timing of the animal’s last meal, the half-life of doxycycline, the concentration of doxycycline needed to suppress gene expression, the possibility that concentrations may be lower in the brain^57^, and the half-life of tTA-protein. Previous studies have suggested that the tagging window opens within 24 hours of food removal^58^ however finer timescale measurements have not been performed to our knowledge. Moreover, we point out that there may be high variability from animal to animal based on individual and strain differences in the rate of doxycycline metabolism and the timing and amount of dox food consumed. Future studies could more precisely measure the onset and duration of doxycycline-dependent gene expression. Alternatively, future studies could shift to more recently developed tagging systems with greater temporal precision.

Other studies have utilized the TRAP2^33,59,60^ system as it labels a smaller proportion of CA1 during a shorter time window, around 5% or less of active cells which is much lower than we observed here (35-40%, <u>Figure 1G</u>, <u>Supplementary Figure 3B</u>). While our results are more closely aligned with opto-tagging approaches in CA1 (∼20%),^11,49^ it is possible labelling a larger pool of mCherry cells contributed to our finding that a subset of engram cells demonstrated the increase in correlation (<u>Figure 4C-D</u>). Given that the amount of CA1 cells labeled with *Fos*-dependent approaches range from 5-40%, this highlights the need for systematic comparison of activity-dependent strategies^61^ with endogenous *Fos* levels, which is both brain-region and experience dependent. While we are not aware of any studies that have described genotype-, dose-, or brain-region specific differences in TRAP2 tagging levels, these methodological differences must be considered when comparing across studies, as experimental decisions such as tamoxifen dosage have influenced the amount of IEG-tagging in other systems.^19^ In addition to the TRAP2 system, newly developed methods that utilize faster timescale light-dependent systems include Cal-Light^62^, and scFLARE/FLiCRE^63–65^ which permit genetic tagging of cells on a seconds-timescale. However, care should be taken when selecting methods as these more temporally restricted tools rely on light activation for tagging that may preclude simultaneous photoexcitation for functional imaging. Thus, employing more temporally restricted techniques combined with experimental designs in which behavior is monitored longitudinally, can be done systematically in future studies to reveal the mechanisms surrounding engram allocation and function.

## Conclusion

Our study reveals fundamental mechanisms of hippocampal engram cell physiology, including how their activity shapes memory allocation and is changed by learning. By combining in vivo imaging, IEG-tagging, behavior and simple spiking models, we concluded that more excitable cells were biased to engram allocation, identified increases in correlated activity after learning and found that cued recall is reflected in both engram activity levels and stable correlation structure. Overall, these findings support a model in which cells are recruited to engram populations based on pre-existing excitability, that learning selectively strengthens their connectivity within their local circuit and that retrieval drives a re-emergence of stabilized coactivity between these cells. Together our results are consistent with a framework in which memory allocation, learning, and retrieval are each reflected in measurable signatures of engram population activity, bridging cellular physiology with systems-level accounts of hippocampal memory.

## Acknowledgements

We would like to thank Jack Giblin, Kivilcim Kilic and the BU Neurophotonics Center for their training and support, and Joseph Zaki for helpful discussion of analyses. Denise Cai and Michael Hasselmo provided valuable comments. AM was supported by an NSF NRT award and an NIH F99/K00 DSPAN award. BBS was supported by a Young Investigator award from the BBRF and by a SEED grant from the BU Neurophotonics Center. CL was supported by a Center for Systems Neuroscience Distinguished Postdoctoral Fellowship. This material is based upon work supported by an NIH Early Independence Award DP5 OD023106-01, NIH Transformative Award, Air Force Office of Scientific Research (AFOSR; FA9550-21-1-0310), the Ludwig Family Foundation, the Pew Scholars Program in the Biomedical Sciences, the Chan-Zuckerberg Initiative, and the Center for Systems Neuroscience and Neurophotonics Center at Boston University.

## Author contributions

Experiment design: AM, SR, BBS

Mouse imaging & behavioral experiments: AM, SC

Data analysis: AM, SC, RS

Spiking model design & simulations: CL, GKO

Manuscript writing: AM

Manuscript editing: AM, CL, BBS, SR, GKO

Declaration of Interest

The authors declare no competing interests.

## Methods

### Key Resources Table

| Reagent or Resource | Source | Identifier |
| --- | --- | --- |
| **Experimental models: Organisms/strains** |  |  |
| C57BL/6J-Tg(Thy1-GCaMP6f)GP5.17Dkim/J | The Jackson Laboratory | Strain 025393 |
| **Bacterial and virus strains** |  |  |
| AAV9-c-fos-tTA-BGHpa | UMass Vector Core | Custom Ordered |
| AAV9-TRE-mCherry | UMass Vector Core | Custom Ordered |
| **Chemicals, peptides and recombinant proteins** |  |  |
| Doxycycline diet (40 mg/kg) | Bio-Serv | Custom ordered dose |
| **Software and algorithms** |  |  |
| Python 3.10 | Python Software Foundation | https://www.python.org/ |
| Arduino | Arduino | https://www.arduino.cc/ |
| CellReg | Sheintuch et al. 2017<sup>66</sup> | https://github.com/zivlab/CellReg |
| CalmAn | Giovanucci et al 2019<sup>67</sup> | https://github.com/flatironinstitute/CalmAn |
| ImageJ (FIJI) | National Institutes of Health | https://fiji.sc/ |
| FreezeFrame4 | Actimetrics, Lafayette, IN | https://actimetrics.com/ |
| Cellpose | Stringer et al., 2021<sup>68</sup> | https://github.com/mouseland/cellpose |

### Experimental Methods

#### Animals & Handling

Transgenic Thy1-GCaMP6f^69^ (C57BL/6J-Tg(Thy1-GCaMP6f)GP5.17Dkim/J, Jackson Labs #025393, 16-24 weeks) were group housed with littermates prior to experiment start and given food and water *ad libitum.* The animal vivarium was maintained on a 12:12-hour light cycle (0700-1900). Following surgery, mice were single-housed to prepare for imaging experiments. All subjects were treated in accord with protocol 201900084 approved by the Institutional Animal Care and Use Committee (IACUC) at Boston University. Prior to imaging experiments, mice were habituated for 5 days to head-fixation.

#### Stereotaxic Surgery & Window Implantation

For all surgeries, mice were initially anesthetized with 3.0% isoflurane inhalation during induction and maintained at 1.5%–2% isoflurane inhalation through stereotaxic nose-cone delivery (oxygen 1L/min). Ophthalmic ointment (Systane) was applied to the eyes to provide adequate lubrication and prevent corneal desiccation. The hair on the scalp above the surgical site was removed using Veet hair removal cream and subsequently cleaned with alternating applications of betadine solution and 70% ethanol. 2.0% lidocaine hydrochloride was topically applied as local analgesia prior to midsagittal incision using an 11 blade scalpel of the scalp skin to expose the skull. A 0.1 mg/kg intraperitoneal (IP) dose of buprenorphine was administered at the beginning of surgery. A unilateral craniotomy was performed with a .5mm drillbit for viral injection. A 10 uL Hamilton syringe with an attached 33-gauge beveled needle was slowly lowered to the coordinates of dorsal CA1: -2.0 AP, +1.5ML -1.5 DV. 400 nL of 1:1 cocktail of AAV9-cFos-tTA/AAV9-TRE-mCherry (UMass Vector Core, titer 1×10^13^ GC/mL) was injected at a rate of 100 nL/min. The needle was allowed to rest at the injection site for 1-2 min prior to and after injection. Incisions were sutured closed using 4/0 Non-absorbable Nylon Monofilament Suture (Matrix Wizard). Following surgery, mice were injected with a 5 mg/kg intraperitoneal (IP) dose of ketoprofen. They were placed in a recovery cage with a heating pad until fully recovered from anesthesia. Histological assessment verified spread of dorsal CA1 viral targeting.

Seven to ten days later, a second surgery was performed to implant a hippocampal window for two-photon imaging. Prior to surgery, dexamethasone (3 mg/kg) was administered IP to aid brain swelling and inflammation. A 3mm craniotomy was created centered at the site of viral injection. Dura was peeled back with a bent 27-gauge needle tip. Cortical tissue washed with sterile saline while aspirated using constant suction attached to 25-30 gauge needle tips. Tissue was aspirated down to white matter and corpus callosum fibers were carefully removed, leaving alveolar fibers intact. Blood was constantly removed via aspiration and absorption with saline-soaked hemostatic collagen sponge (Goodwill Hemosponge Absorbable Gelatin Sponge).

For the window implant, a 3mm diameter, 1.5mm deep stainless steel circular cannula (Ziggy’s Tubes & Wires) was affixed to a 3 mm glass coverslip (#1 thickness, Thomas Scientific 1217N66) using UV curable optical adhesive (Norland). The cannula window was lowered carefully into the craniotomy site and Vetbond was used to adhere the edges of the cannula to the skull. The scalpel and drillbit was used to hatch the skull surface for best adhesion to the headplate. A custom stainless steel kinematic headplate^21^ was affixed to the skull with the cannula centered using Metabond (Parkell). Mice were allowed to recover at least 1 week prior to handling & imaging took place 2 weeks after surgery.

#### Two-photon Imaging

All imaging was performed on a Bruker Ultima Investigator in Prairie View (v5.4). Excitation was delivered with a tunable Ti:Sapphire laser (Insight X3, Spectra Physics) at wavelengths of 920nm or 1040 nm and a primary dichroic (700 SP) split excitation and fluorescence light and a 770 SP IR blocker ensured no laser power reached the PMTs. On each imaging day, image series were collected at a frame rate of 30Hz with resonant galvos and resolution of 512×512 pixels. All imaging was with a 16x Nikon (3.0 mm, 0.8 NA) water immersion objective. Laser power was controlled with an electro-optic modulator (Conoptics 305-105-20) and post objective power ranged from 10-80 mW and did not exceed 100 mW. Emitted fluorescence was split into red and green channels with an emission filter cube set (565 LP dichroic, 525/70 for the green channel and 595/50 for red) and detected with two PMTs, for green (Hammamatsu H10770PB-40) and red (Hammamatsu H10770PB-50) channels. For recording spontaneous activity, 1-2 FOVs were recorded per animal for 22 minutes on Day 0 and Day 4 at 920nm. 1 FOV was excluded from 5 out of 12 mice due to either unreliable cell registration or resonant galvo image artifacts. For recording dual color GCaMP/TRE-mCherry expression, the same FOVs were recorded for ∼1 minute (2000 frames) at a wavelength of 1040 nm. All dual color data was recorded at identical power settings on Day 0 and Day 4 in order to characterize mCherry fluorescence change across days.

During imaging mice were placed on a custom rotating disk that enabled them to locomote. A kinematic headplate and mount system was custom designed to ensure reliable registration of imaging FOV day to day and allowed for locomotion.^20^ The imaging apparatus consisted of the kinematic mount held in place with ½” Thorlabs posts above a horizontal running disk. The rotating acrylic disk was 7” in diameter and a soft rubber adhesive was used to provide the mice traction for running on the disk. The center of the disk was screwed into an adjustable collar (McMaster Carr 9410T1) and mounted on a rotary encoder (US Digital H1-5000-IE-D) to measure running. A custom arduino script was used to record and synchronize image acquisition with running data from the disk. Wheel data was recorded in 13 of 16 mice for <u>Figures 1</u>-<u>4</u> (CFC/HC mice) and for all mice in <u>Figure 5</u> (TFC mice).

#### Running behavior

To collect locomotion data, an Arduino script was used to record timestamped positional data from the mouse’s running disk. Timestamps were aligned to imaging frames by interpolating wheel positional data with known frame times from Prairie View xml metadata files. To determine walking/running epochs, a velocity threshold of 1 cm/s was imposed to select for frames to account for spurious movement of the running disk. Run epochs were selected requiring that the animal be greater than one for longer than 0.5 seconds and rest epochs were selected requiring the animal to be still for at least 0.5 seconds.

#### Fear Conditioning Tagging and Behavior

Prior to surgeries, mice were placed on a special diet of doxycycline chow (40 mg/kg, BioServ). When dox chow was removed (end of D0), this opened the window for activity-dependent viral labeling during the tagging period. For tagging, HC mice remained untouched in the home cages, while FC mice were placed in a conditioning chamber with plexiglass walls and a grid floor (17.78 cm L × 17.78 cm W × 30.48 cm H; Coulbourn Instruments) for a 7-minute conditioning session. Grid floors were connected to precision shockers and delivered a series of three footshocks (2 s, 1.0 mA intensity) at times 200s, 280s and 360s. Immediately after fear conditioning, the doxycycline diet was replaced in all mice cages immediately after tagging and mice remained on doxycycline diet for the duration of the imaging protocol (until after Day 4). Dox was removed after the conclusion of imaging sessions to allow full expression of tet-driven mCherry and check for injection spread (data not shown). For fear conditioning, video data were collected via overhead cameras (Computar) that interface with FreezeFrame (Actimetrics). FreezeFrame was used to control delivery of the foot shocks and perform freezing analyses. Freezing during fear conditioning was defined as bouts of 1.25 s or longer with minimal changes in pixel luminance.

#### Trace Fear Conditioning

Auditory trace fear conditioning took place on Day 2. After imaging on Day 0, doxycycline was replaced with standard chow and mice were left undisturbed until Day 2. Trace conditioning was performed in a Med Associates Chamber with a nine-minute protocol in MedPC. Three auditory stimuli (2.8 kHz, 20 s, pure tone) were presented at 180s, 300s, and 420s. 20 seconds after each tone termination a 2 second, 1.0 mA shock occurred.

#### Imaging during auditory CS presentation

Recording was performed similar as described above, where 27 minutes of spontaneous baseline activity was recorded on Day 0. Two days after trace fear conditioning, the same field of view was recorded again. First, 10 minutes of spontaneous activity was recorded before ten 20-second presentations of the CS+ (2.8kz pure tone) was played, followed by an 80 second ITI. In 5 mice, CS-trials were randomly interleaved (5 trials of each type) on Day 5 of imaging. For CS-trials, Bpod delivered a 12kHz pulsed (1Hz, 50% duty cycle) tone for 20s.Tone stimuli were delivered via a speaker and Teensy (Arduino) controlled by BPod (Sanworks) and synchronized with imaging frames. During tone presentations PrairieView recorded TTL pulses sent from BPod to align CS and imaging frames.

#### Immunohistochemistry

For histological section experiments (<u>Supplementary Figure 3</u>), mice were euthanized 90 minutes after behavior with 3% isoflurane and transcardially perfused with cold phosphate buffered saline (PBS), followed by 4% paraformaldehyde (PFA) in PBS. Brains were extracted and stored in 4% PFA at 4℃ for at least 24 hrs. Brains were sliced coronally at 40um on a vibratome and stored in 0.01% sodium azide. Staining was performed to amplify mCherry and endogenous cFos. Slices were selected and washed with PBS at room temperature for 10 minutes three times. Tissue was then blocked in 5% bovine albumin serum (BSA) at room temperature for 2 hrs. Slices were then placed in solution of 1% BSA with primary anti-bodies (1:5000 guinea pig anti-RFP [Synaptic Systems 390 004] and 1:1000 Rabbit anti-cfos [Abcam ab190289]) and left to incubate for 48 hrs. After the incubation period, tissue was washed in phosphate-buffered saline with triton (PBST) for 10 minutes three times. The slices were then stained with secondary antibodies (1:500 Goat anti-Guinea 555 [Thermofisher A-21435] and 1:200 Goat anti-Rabbit 488 [Thermofisher A-11008]) and left to incubate for 2 hours at room temperature. Tissue was then washed in PBS at room temperature for 10 minutes three times, then mounted on slides with Vectashield HardSet Mounting Medium with DAPI (Vector Laboratories H-1500-10) and coverslipped.

#### Confocal Imaging and Cell Counts

Coronal brain sections were imaged using a Zeiss LSM 800 epifluorescence/confocal microscope equipped with a 20x/0.8 NA objective using the Zen2.3 software. Images of dorsal CA1 (dCA1) were acquired in DAPI (405nm), RFP (555 nm), and cFos (GFP, 488nm) channels. For each animal, 3-5 slices were imaged and averaged for analysis.

Confocal images of dCA1 were preprocessed in FIJI (ImageJ). A custom Fiji macro was used to split channels, adjust brightness and contrast, and separate z-stacks. Regions of interest were delineated with the polygon tool to isolate the pyramidal cell layer of dCA1. For each z-stack, the three central slices with the highest signal intensity were selected for quantification. Cell segmentation and counting were performed using a custom-trained Cellpose3^68^ model and resulting cell counts were exported as NumPy files for averaging across slices, animals, and experimental groups.

In cases where DAPI counts could not be obtained directly, values were estimated on the relationship between countable DAPI cells and the measured area of the DAPI-labeled region. The percentage of mcherry-positive cells was calculated as (mCherry/DAPI x100). For each mouse, a single averaged value across all slices was generated and used for subsequent statistical analyses.

### Data Analysis

#### Image Processing and ROI Extraction

Images were first spatially resized in Fiji (method: averaging, bilinear interpolation) to 256×256 (1.58 um/pixel resolution). All image processing was done using custom Python scripts using the CaImAn python package.^67^ Images were first motion corrected using NoRMCorre with non-rigid motion correction implemented in CaImAn. Day 4 images were registered to the average template image from Day 0 so cellular ROIs to allow for in alignment across days. Similarly, two-color images (1040 nm time series) were aligned to the 920nm functional images as follows. 1040 nm green channel images were motion corrected to the 920 nm GCaMP average image template and these shifts were applied to the 1040 nm red channel images in order to align green/red channels with the ROIs extracted on the 920 nm images. In some cases, the red channel images were motion corrected to the Day 0 average red channel image template and shifts were applied to the green channel. After motion correction, CNMF was used to extract cellular ROIs and dF/F traces. Registration of cell ROIs across days was performed with the CellReg MATLAB GUI.^21^ Visual inspection of spatial ROIs and cell registration was performed, and inaccurate spatial ROIs were excluded. For this step, first CellReg output was visualized as a whole with color coded ROIs for registered cells (<u>Supplementary Figure 2A</u>). Next, each ROI pair was plotted on top of the session’s mean template image and correlation image (calculated with CaImAn’s ‘*caiman.summary_images.local_correlations*’ function) and manually verified as a true registered cell pair (<u>Supplementary Figure 2B</u>). Additionally, ROIs with a SNR less than 4 were discarded based on visual inspection of lower SNR traces. We did this by thresholding ROIs using CaImAns SNR metric for each ROI from ‘*cnmf.estimates.SNR_comp*’. Cell ROIs were then manually classified as engram or non-engram based on user-picked threshold of mCherry intensity and visual evaluation of their mCherry expression on Day 4, taking into account the soma intensity relative to the local background of each cell (<u>Figure 1D</u>, see mCherry expression analyses below).

#### mCherry expression analyses

We analyzed mCherry expression using motion-corrected, averaged red channel images acquired at 1040 nm following the two-color registration procedure described earlier. A robust increase in overall mCherry fluorescence was observed from Day 0 (D0) to Day 4 (D4) (<u>Supplementary Figure 1B-C</u>), indicating widespread mCherry labeling during the Dox withdrawal period. Notably, this increase was evident across the entire cell population (<u>Supplementary Figure 1B</u>).

To identify engram cells, we established a threshold for mCherry pixel intensity on D4. These thresholds were manually evaluated to accommodate field-of-view (FOV) signal heterogeneity and to minimize false positives or negatives that could arise from applying a uniform threshold across FOVs with non-uniform luminance. To validate our approach, we quantified mCherry intensity relative to each cell’s local background. This was achieved by scaling each cell’s ROI coordinates by a factor of 1.5 around its center of mass to generate a larger surrounding ROI. The background region was isolated by subtracting the original ROI from this expanded ROI, and the mean intensity of this surrounding area was calculated. The mean soma intensity was then divided by this background intensity to yield the Relative Fluorescence metric (<u>Figure 1D</u>). Engram cells displayed higher mCherry fluorescence relative to their local backgrounds (<u>Figure 1D</u>) and also exhibited elevated raw mCherry intensities (<u>Supplementary Figure 1C-D</u>). We further assessed changes in mCherry expression by calculating the fold change from D0 to D4 (<u>Supplementary Figure 1F-G</u>). For each cell, this was computed by dividing the D4 mean ROI intensity by the D0 intensity.

Finally, to control for potential leaky expression from incomplete doxycycline suppression of TetTag constructs, we excluded cells with high mCherry expression at D0 from the engram classification. Specifically, we removed cells with expression levels exceeding 2.5 standard deviations above the mean ROI intensity at D0. These cells accounted for approximately 2% of the active CA1 population (<u>Supplementary Figure 1E</u>).

#### Calcium deconvolution and event rates

All dF/F traces underwent calcium deconvolution to infer discrete event times. For event detection we utilized an L0 penalty method described in References 59-60.^70,71^ Briefly, each cell’s dF/F trace is modeled as a noisy readout of underlying calcium fluorescence dynamics. Two parameters are used to fit event times to each cell’s trace: gamma and lambda. Gamma is determined by the half-life of the indicator which we set to 0.14 s as reported for the Thy1-GCaMP6f GP.5.17 line.^69^ Lambda controls the strictness of the L0 penalty and therefore the sparseness of events, with higher lambda values resulting in higher sparsity. We estimated lambda for each cell similarly to previous reports^72^ and adjusted the minimum event size to 5 standard deviations of the noise in the dF/F of each cell. This threshold was chosen based on manual inspection of deconvolved traces with lower SNR and sparser activity. Mean event rates were calculated using the weighted sum of deconvolved event sizes, in order to account for calcium transients with larger amplitude as follows. For each cell, all detected calcium events had an associated non-zero magnitude, which were summed over the course of each imaging session and divided by total session time. This ensured smaller transients were not treated the same as larger transients when determining calcium event rate. For trace conditioned mice, deconvolved event rates were calculated during each 20s CS trial and then z-scored to the baseline period (<u>Figure 5E</u>).

#### Correlation analyses

Deconvolved traces for cells were binned using overlapping bins of 4, 8,16,25, 50, 80, 100, and 125 frames. Pairwise Spearman correlations between every pair of cells simultaneously recorded were performed. Only cells active in both Day 0 and Day 4 sessions were used for this analysis. Pairwise correlations were grouped by cell identity and the mean correlation coefficient for all engram/engram, engram/non-engram or non-engram/non-engram was calculated for each animal <u>Figure 4A</u>. When calculating mean correlation during rest and run states (<u>Figure 4B-C</u>), velocity was thresholded as described above, rest and run epochs required a minimum duration of 1s to be included, and sessions were matched for duration in each state. For thresholding correlated pairs, we chose all pairwise correlations exceeding p-value thresholds ranging from 0.01 to 10e-5 and looked at positively correlated pairs. We took the mean of these thresholded correlations within each population, animal and day and calculated the normalized change as D4/D0.

For <u>Figure 5F</u> we repeated this approach but calculated the change in Spearman correlation for each pair of cells from pre-CS+ to CS+ presentations. For each cell-pair, the Spearman coefficient was calculated for the pre-CS+ activity and CS+/trace period activity (concatenated across trials), and correlations with *p<*0.05 were kept. The difference in Spearman’s rho across these two periods reflected the change in correlation induced by the CS+. This difference was z-scored across all pairwise comparisons within each animal and the absolute value (both increases and decreases) was plotted in <u>Figure 5F</u> for different bin sizes of the original activity.

#### Geometric distance

To compare the similarity of the correlation matrices across days (<u>Figure 4B</u>) or CS+ presentation (<u>Figure 5G</u>), we employed a log-Euclidean Riemannian metric from Ref 27. This metric is used to compare geodesic differences between two correlation matrices. These correlation matrices were generated by Spearman correlations across all pairs of cells’ deconvolved activity traces (not binned). For two correlation matrices A and B the distance metric was:

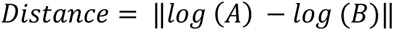

Where log() is the matrix logarithm. To account for differences in population sizes (engram versus non-engram) this distance was normalized by the square root of the population size.

#### Rastermap and sequence analysis

To identify sequential dynamics in the deconvolved activity, event traces were binarized and given to rastermap. Rastermap was run using the following parameters {n_clusters=None, nPCs=10, time_lag_window=60, locality=0.1, time_bin=15}, which were similar to recommended parameters for single-cell sorting in hippocampal navigation data. For <u>Figure 3E</u>, cells that participated in rastermap-sorted sequences were chosen from sorted rastermaps, as their sorting algorithm groups sequentially active cells together (<u>Figure 3D</u>). For <u>Figure 3F</u>, velocity correlation was calculated as the Pearson correlation between the summed activity of the population (engram or non-engram) and velocity.

#### Hierarchical Bayesian GLMs

These HB-GLMs leveraged partial pooling of event rates across animals to increase statistical power. In these HB-GLMs, the slope coefficient *β*_1_ represents the effect of a cell’s event rate on the likelihood of tagging, and the intercept *β*_0_ captures baseline tagging variability. Both parameters were estimated for each animal individually under a group-level prior distribution. To avoid prior bias, we used noninformative priors centered *β*_0_= 0 and *β*_1_= 0, corresponding to a null hypothesis of no relationship between activity rate and tagging likelihood (<u>Supplementary Figure 4D</u>).

We implemented four variants of a hierarchical Bayesian logistic regression model to analyze the relationship between event rates and population outcomes (i.e., were cells tagged or not during learning) across different animals. Models were fit using the python packages arviz and pymc. Separate models were fit for event rates during 1) rest periods only (Rest model), 2) running bouts only (Run model), 3) both resting and running, with separate regressors for cell’s event rates during rest and run (Combined model), and 4) a null model (*β*_1_ = 0) (<u>Supplementary Figure 4A,D</u>).

For each animal i and cell j:

Where s is the sigmoid function:

The linear predictor varied by model:

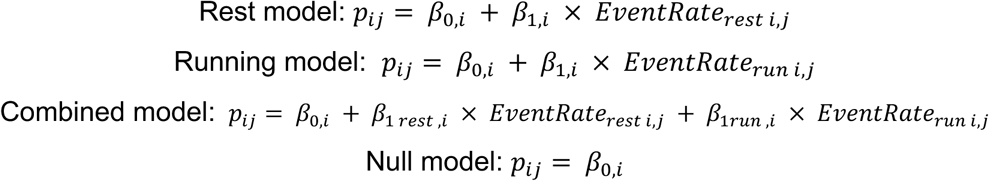

where the individual-level parameters follow:

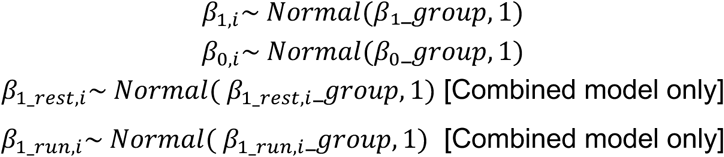

And the group-level parameters have the following prior distributions:

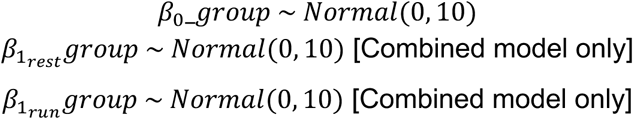

To evaluate model performance, we compared the expected log predictive density using leave-one-out cross-validation (ELPD-LOO). The combined model showed the best predictive performance, with the highest ELPD-LOO value (<u>Supplementary Figure 4D</u>, Combined: -2438). In contrast, both the run-only and null models underperformed relative to the rest-only and combined models, as reflected in their lower LOO values (<u>Supplementary Figure 4D</u>, run: -2445, null: -2456). The Combined model performed significantly better than the Null model (ELPD difference: 17.6 ± 7.2). Notably, the Rest-only model was nearly as informative as the Combined model, with only a small, non-significant ELPD difference, suggesting that rest activity alone captured most of the predictive structure (<u>Supplementary Figure 4D</u>, ELPD-LOO combined: - 2438, rest: -2443). These results are consistent with our prior linear regression analyses (<u>Figure 2</u>), as models incorporating rest-period activity demonstrated improved predictive performance. For both Combined and Rest models, the estimated *β*_1 *rest*_ values for individual animals were consistently positive and greater than the *β*_1 *run*_ values in both combined and rest-only models (<u>Supplementary Figure 4B-C</u>). Finally, group-level posterior distributions reinforced these effects, as the 95% high-density intervals (HDIs) for *β*_1 *rest*_excluded zero for rest and combined models (<u>Supplementary Figure 4C</u> Combined *β*_1 *rest*_ [0.63, 2.39], Rest *β*_1_ [0.85, 2.56]), while the *β*_1 *run*_ coefficients were less reliable and had weaker explanatory power (Run *β*_1 *run*_: [0.06, 1.37], Combined *β* _1 *run*_ [-0.08,1,23]) Taken together, these comparisons suggest that rest activity alone captures most of the predictive structure, while adding run activity provides limited additional value.

#### Histology Confocal Imaging and Cell Counts

Coronal brain sections were imaged using a Zeiss LSM 800 epifluorescence/confocal microscope equipped with a 20x/0.8 NA objective using the Zen2.3 software. Images of dorsal CA1 (dCA1) were acquired in DAPI (405nm), RFP (555 nm), and cFos (GFP, 488) channels. For each animal, 3-5 slices were imaged and averaged for analysis.

Confocal images of dCA1 were preprocessed in FIJI (ImageJ). A custom Fiji macro was used to split channels, adjust brightness and contrast, and separate z-stacks. Regions of interest were delineated with the polygon tool to isolate the pyramidal cell layer of dCA1. For each z-stack, the three central slices with the highest signal intensity were selected for quantification. Cell segmentation and counting were performed using a custom-trained Cellpose3 model and resulting cell counts were exported as NumPy files for averaging across slices, animals, and experimental groups.

In cases where DAPI counts could not be obtained directly, values were estimated on the relationship between countable DAPI cells and the measured area of the DAPI-labeled region. The percentage of mcherry-positive cells was calculated as (mCherry/DAPI x100). For each mouse, a single averaged value across all slices was generated and used for subsequent statistical analyses.

#### Statistical analysis

All statistical analyses were performed in Python using the pingouin (v0.5.3) or statsmodels (v0.13.5) packages. We assessed normality using the Shapiro-Wilk test to guide the choice of statistical tests. For comparisons across groups and timepoints, we employed either 2-Way Repeated Measures ANOVAs (for balanced, parametric data) or linear mixed effects models (for unbalanced, non-parametric, or repeated-measures designs), with random intercepts for mice or FOVs when applicable. Post hoc comparisons were conducted using paired (across mice or FOVs) or unpaired t-tests for parametric data, and Wilcoxon signed-rank or Mann-Whitney U tests for non-parametric data, with multiple comparisons corrected using Benjamini-Hochberg FDR, alpha=0.05. For post-hoc contrasts within linear mixed effects models, we used Wald contrast tests, in which a linear combination of fixed-effect coefficients was tested against zero using a z-statistic derived from the model’s estimated covariance matrix (two-tailed). Data are presented as mean ± SEM unless otherwise indicated. For regression analyses, 95% confidence intervals are reported.

### Spiking Network Model of CA3-CA1

#### Spiking Network Model of CA3-CA1

We model each neuron’s spike train as a Poisson process with time dependent rate *r*_*i*_(*t*) given by

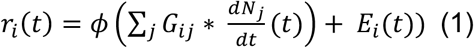

Here, *N*_j_(*t*) counts the total number of spikes neuron *j* has fired up to time *t*, and its derivative 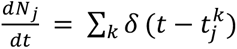 describes the neuron’s spike train as a sum of Dirac delta functions. *G* is a matrix of synaptic interaction filters, and ∗ denotes a convolution, i.e. 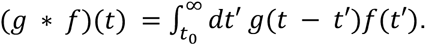. We take 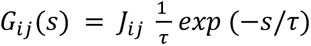 (for *s* ≥ 0) where *J*_*i*j_ gives the synaptic weight from neuron *j* onto neuron *i*. We take *ϕ* to be threshold-quadratic, i.e. *ϕ*(*x*) = max(*x*, 0)^2^.

Our synaptic weight matrices were constructed to reflect recurrent E-E connectivity in CA3, as well as recurrent E-I connectivity in both CA1 and CA3 and feedforward connectivity from CA3 to CA1 (<u>Figure 5A</u>). We base our weight matrix *J* on the connectivity of regions CA1 and CA3 of the hippocampus. We further subdivide both CA1 and CA3 into excitatory engram neurons (E), excitatory non-engram neurons (N), and inhibitory neurons (I). The regional connectivity is given by a 6×6 matrix *J̅*.

Our synaptic weight matrices were constructed to reflect recurrent E-E connectivity in CA3, as well as recurrent E-I connectivity in both CA1 and CA3 and feedforward connectivity from CA3 to CA1 (<u>Figure 5A</u>). We base our weight matrix *J* on the connectivity of regions CA1 and CA3 of the hippocampus. We further subdivide both CA1 and CA3 into excitatory engram neurons (E), excitatory non-engram neurons (N), and inhibitory neurons (I). The regional connectivity is given by a 6×6 matrix <u>*J*</u>.

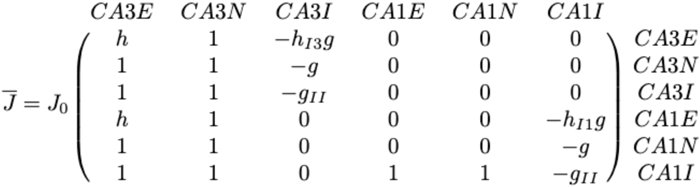

Here, the parameter ℎ describes the strength of engram to engram synapses (in CA3 and CA3 to CA1), relative to the baseline strength of excitatory to excitatory synapses, the parameter *g* describes the relative strength of inhibitory to excitatory connections, the parameter *g*_*II*_ describes the relative strength of inhibitory to inhibitory synapses, and the parameters *h_I3_* and *h_I1_* describe the strength of the homeostatic inhibitory plasticity, i.e. the increased inhibition recruited by engram cells, in CA3 and CA1, respectively.

Neuron to neuron connectivity is described by a block-wise Erdős-Rényi adjacency matrix *A*, with sparse excitatory connectivity and dense inhibitory connectivity given by *p*_*EE*_ = 0.1, *p*_*IE*_ = 0.8, *p*_*II*_ = 0.8, *p*_*EI*_ = 0.8. The connection probabilities do not differ between engram and non-engram cells. Finally, the neuron-to-to neuron weights *J*_*i*j_ are given by 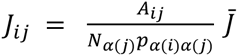 _*α*(*i*) *α*(*j*)_ where *α*(*i*) gives the region containing neuron *i*.

#### Estimating population rates and correlations

In addition to using our stochastic spiking model to simulate neural activity, we calculate estimates for the expected firing rate *r*_*i*_ of each neuron as well as the expected correlation *C*_*i*j_ between each pair of neurons. For more details, see Refs ^73–76^. To find a self-consistent equilibrium rate *r*_*i*_ for each neuron, we substitute *r*_j_ for 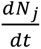 in Equation 1 and replace convolution with *G*_*i*j_ with multiplication by *J*_*i*j_ because the expected rate *r*_j_ is constant. We then find *r* by numerically solving

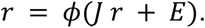

In addition to finding the individual expected rate for each neuron for a fixed neuron-level connectivity matrix *J*, we can find the expected rate of a neuron in each population, where the expectation is taken both over the random connectivity matrix *J* and the stochastic spiking. Letting <u>*r*</u> denote the vector of expected rates for each population and <u>*E*</u> denote the mean input to each population, we have

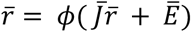

We again solve this numerically to find <u>*r*</u>. Note that these estimates are all for the mean-field rate, i.e. they do not include corrections to the firing rate resulting from correlations in the input interacting with the nonlinear function *ϕ*. To account for this, we add a one-loop correction term.^64^

To estimate the expected spike train covariance matrix Z, we linearize around the expected rate and apply the formula

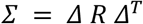

where *Δ* = (*I* − *ϕ*′(*r*) *J*)^−1^, *R* is a diagonal matrix with *R*_*ii*_ = *r*_*i*_, and *ϕ*′(*r*) is a diagonal matrix with *ϕ*′(*r*)_*ii*_ = *ϕ*′(*r*_*i*_). We then compute the correlation matrix *C* from the covariance matrix Z by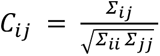.

As with the rate, we also estimate the expected value for the population-level correlation, i.e. the matrix *C* with entries *C*_*αβ*_ giving the expected correlation between a neuron in population *α* and a neuron in population *β* where the expectation is taken over both the random connectivity matrix *J* and the stochastic spiking. Letting Σ_*αβ*_ denote our estimate of the expected covariance between a neuron in population *α* and a (distinct) neuron in population *β*, we have

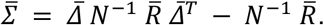

Here, we define 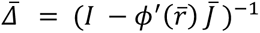 and use *N* to denote the diagonal matrix with *N*_*αα*_ giving the number of neurons in population *α*.

For each population, we can also compute the expected variance of a neuron, which is given by 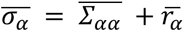. Finally, we compute the population correlation matrix 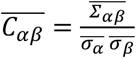.

## SUPPLEMENTARY FIGURES

**Supplementary Figure 1.**
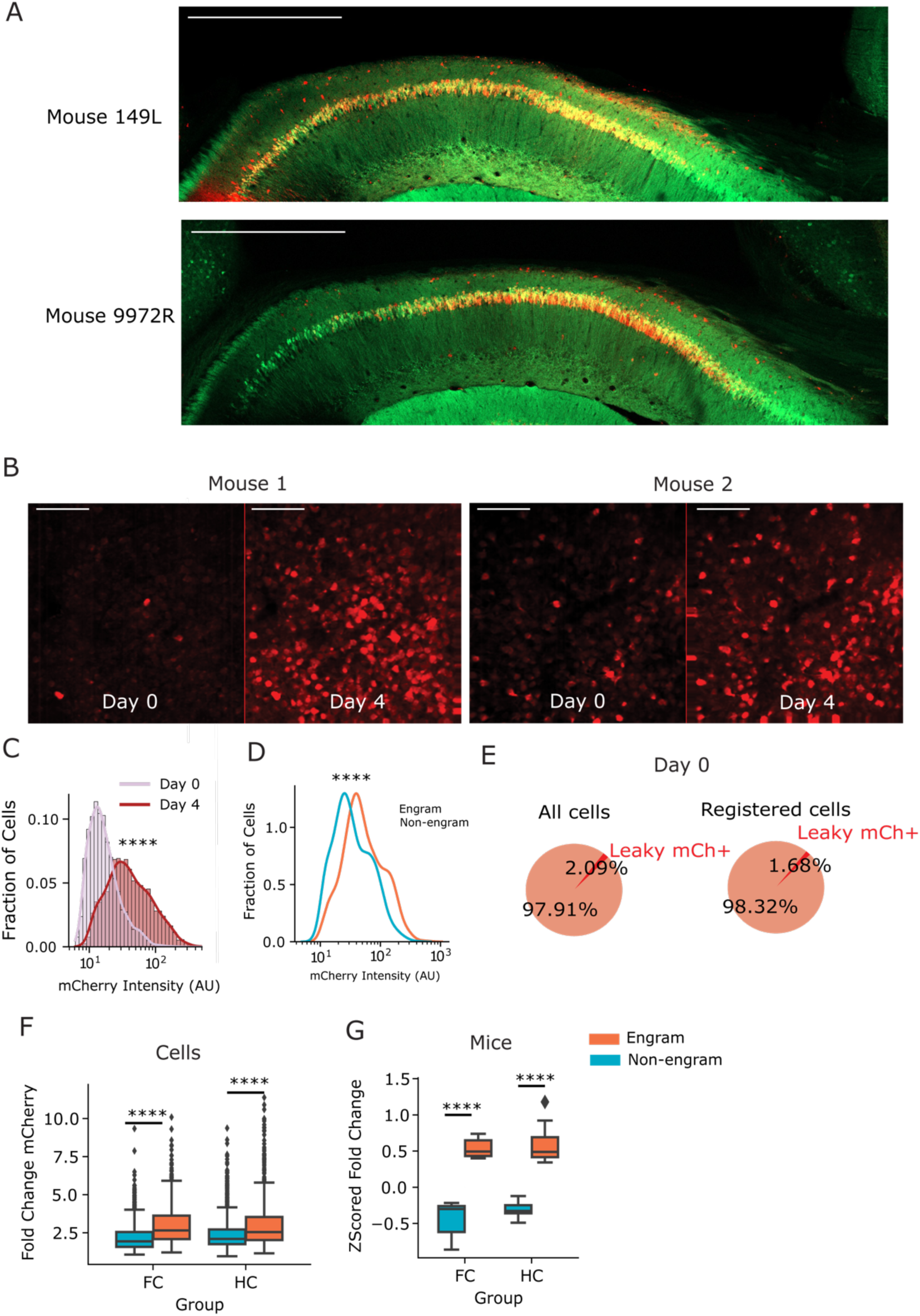
Characterization of tetTag with in vivo imaging & fear conditioning behavior. A) Two example histological sections of FC imaging mice. Mice were taken Off dox at the end of the experiment to check viral spread. Scale bar is 200 μm. B) Two example FOVs for tet-driven mCherry expression across days in vivo. mCherry tagging increases after Day 2 and is measured on Day 4. Scale bars are 100 μm. C) Increase in mCherry intensity of cells quantified from Day 0 and Day 4. Wilcoxon signed-rank test W=119, p=0.0) D) On Day 4, tagged engram cells had significantly higher expression of mCherry (Mann-Whitney U Test: U= 952681.0, p=4.1e-36) E) Percentage of active cells on Day 0 with high, leaky mCherry expression. Approximately 2% of active cells demonstrated high expression on Day 0 before tagging and were not classified as engram cells. F) Engram cells in both FC and HC mice demonstrated a higher Fold Change in mCherry expression from Day 0 to Day 4 (FC Mann Whitney U Test U= 101666.0, p<0.0001, HC U= 143761.0, p<0.0001) G) Mean Z-Scored Fold Change in mCherry expression across mice. 2Way RM ANOVA Group X Population (Group F 1.639100, p= 0.2; Population F=161, p=3.9e-13; Interaction F = 0.18, p=0.6

**Supplementary Figure 2.**
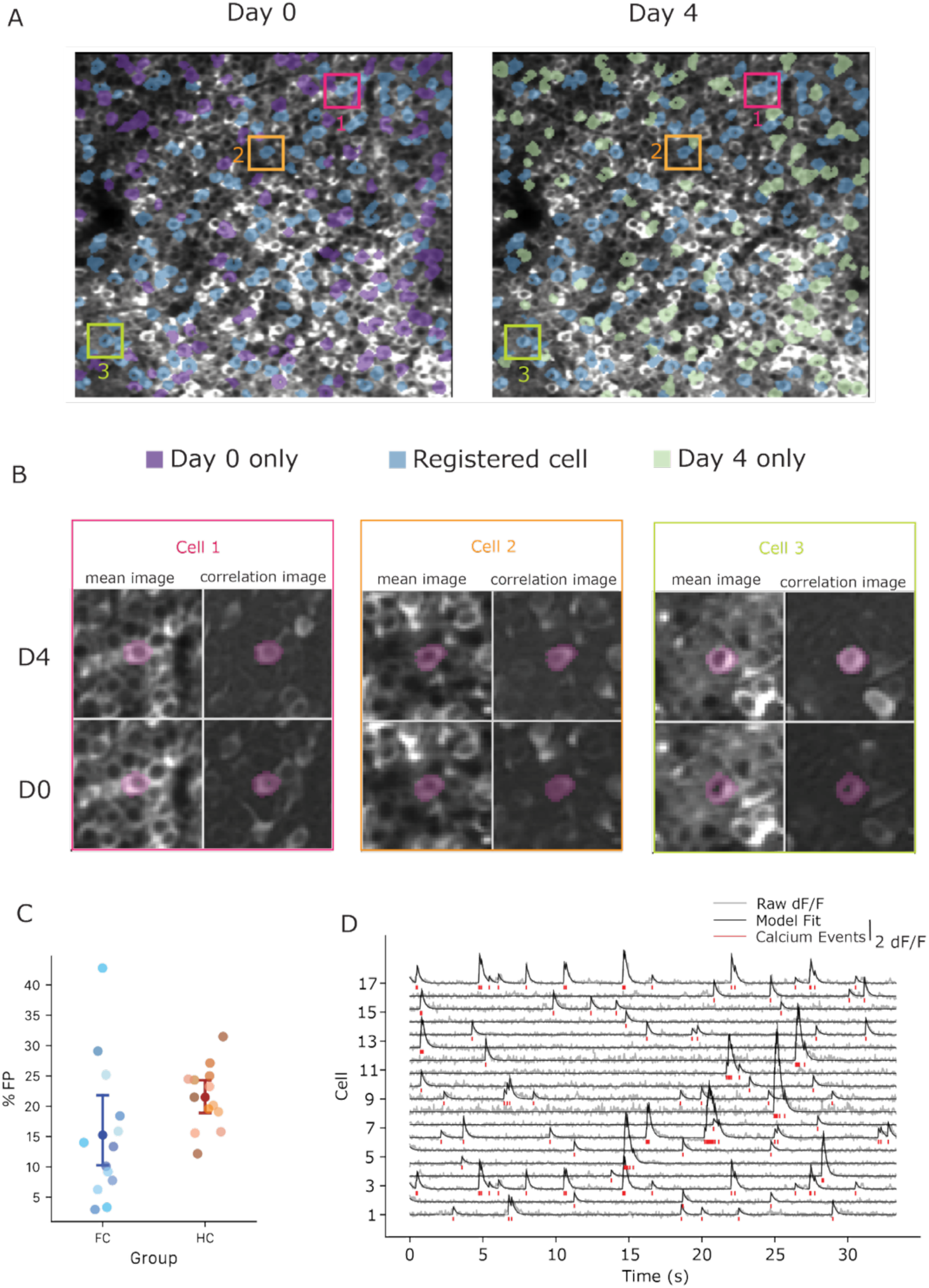
Cell registration quantification and calcium deconvolution. A) Example FOV of cell registration across 4 days. Blue cells: registered, green cells: D4 only, purple cells: D0 only B) Example registered cell pairs plotted over mean projection image and correlation image on each day C) Proportion of False Positive (FP) registered cell ROIs for each mouse/FOV. Manual evaluation of cells considered false positives if 1 of 3 criteria were met: 1) CNMF spatial footprint did not represent a whole/true cell, 2) if the identified pair from CellReg was not a true match or 3) if CNMF spatial footprint overlapped between 2 cells or 4) the CNMF spatial footprint was in a dark area of the FOV. D) Example dF/F traces with respective deconvolved events below. Deconvolution model fit (black) overlaid on top of raw dF/F (gray), calcium events shown below each transient (red).

**Supplementary Figure 3.**
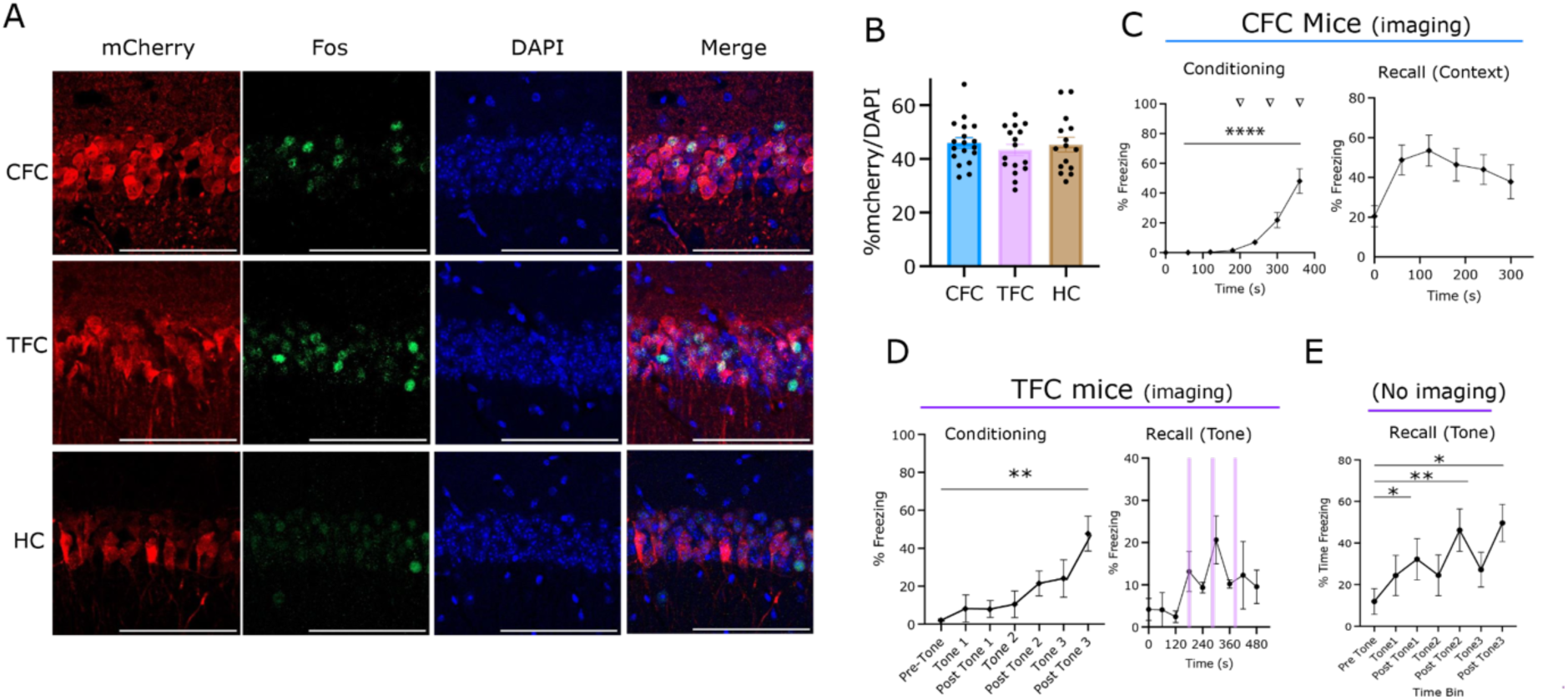
Tet-tag histology and fear conditioning behavior. A) Representative histological sections of tet Tag (cFos-tTA, TRE-mCherry) expression in CA1. 3 groups of mice CFC (n=18), TFC (n=15) or HC (n=16) underwent tagging, with either contextual fear conditioning (CFC), trace fear conditioning (TFC) or remained in homecage (HC) on Day 2 of a 48 hour off-dox window. 2 days after tagging, mice underwent recall (or remained in homecage) and were sacrificed 90 minutes later for Fos expression. Scale bars are 100 microns. B) Percentage of mCherry / DAPI (left) tagged cells or Fos (right) in each group. One Way ANOVA, *p=*0.70. C) Freezing behavior of CFC mice from 2p experiments. Freezing during CFC (left, One Way ANOVA p<0.0001) and Recall (right). Recall took place 2 days after imaging ended (Day 6 relative to first imaging day). D) Freezing behavior of TFC mice from 2p experiments. Freezing during TFC (left, One Way ANOVA p=0.0004 n=8) and Tone recall (right). In a subset of mice (n=3) Recall took place 2 days after imaging ended, in a neutral context (Day 6 relative to first imaging day). E) Additional cohort of TFC mice that demonstrate freezing to tone CS in a neutral context. One Way ANOVA, pairwise t-tests. * p<0.05, ** p<0.01.

**Supplementary Figure 4.**
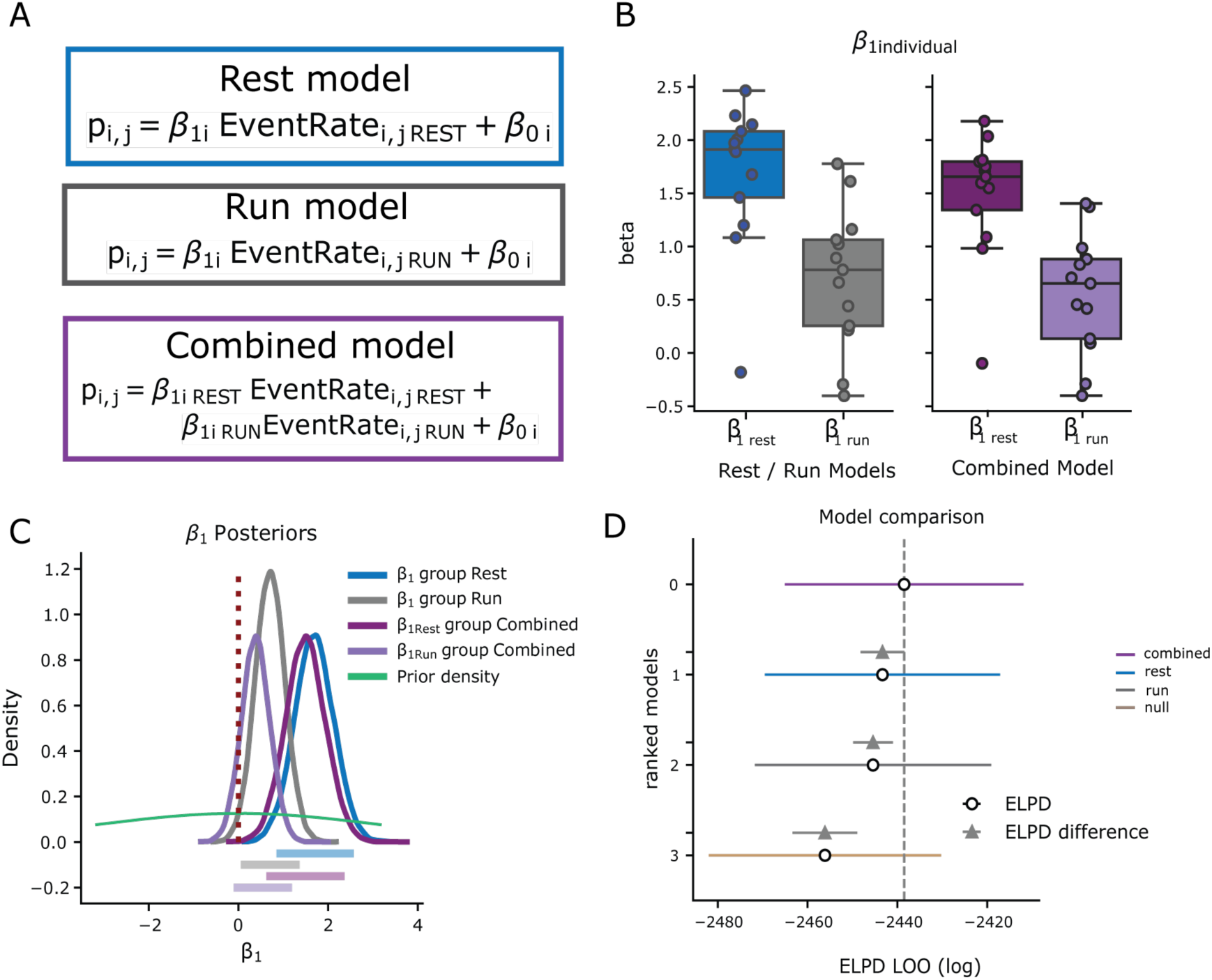
Model-based analysis of engram cell firing. A) Setup for different variants of hierarchical GLM. Null model not shown, but equivalent to *ß_1_* = 0 (intercept only). For mouse i, and cell j, probability of Fos-tagging is modeled as a function calcium event rate governed by animal level parameter *ß_1_*, and *ß_0_* represents a bias term in baseline tagging variability across mice. B) Individual *ß_1_* values for each animal across rest, run and combined models. *ß_1_* in the rest only model (blue) and *ß_1_* ^rest^ in the combined model (dark purple) both have greater magnitudes across all individual animals. C) Posterior distributions for group level *ß_1_* _group_ coefficients compared to prior distribution (green). Horizontal lines indicate 95% high-density interval for each model group level posterior. *ß_1_* for rest model (blue) *ß_1_* ^rest^ (dark purple) both span intervals further from 0 indicating event rate during rest is more informative for predicting likelihood of tagging. D) Model comparison with ELPD-LOO (Expected Log Predictive Density from Leave-One-Out Cross-Validation) metrics for each model. Higher value indicates better predictive power. Standard error of difference between models shown in gray triangle intervals. The combined model performs the best, while run and null models perform significantly worse. The rest only model performs close to the combined model.

**Supplementary Figure 5.**
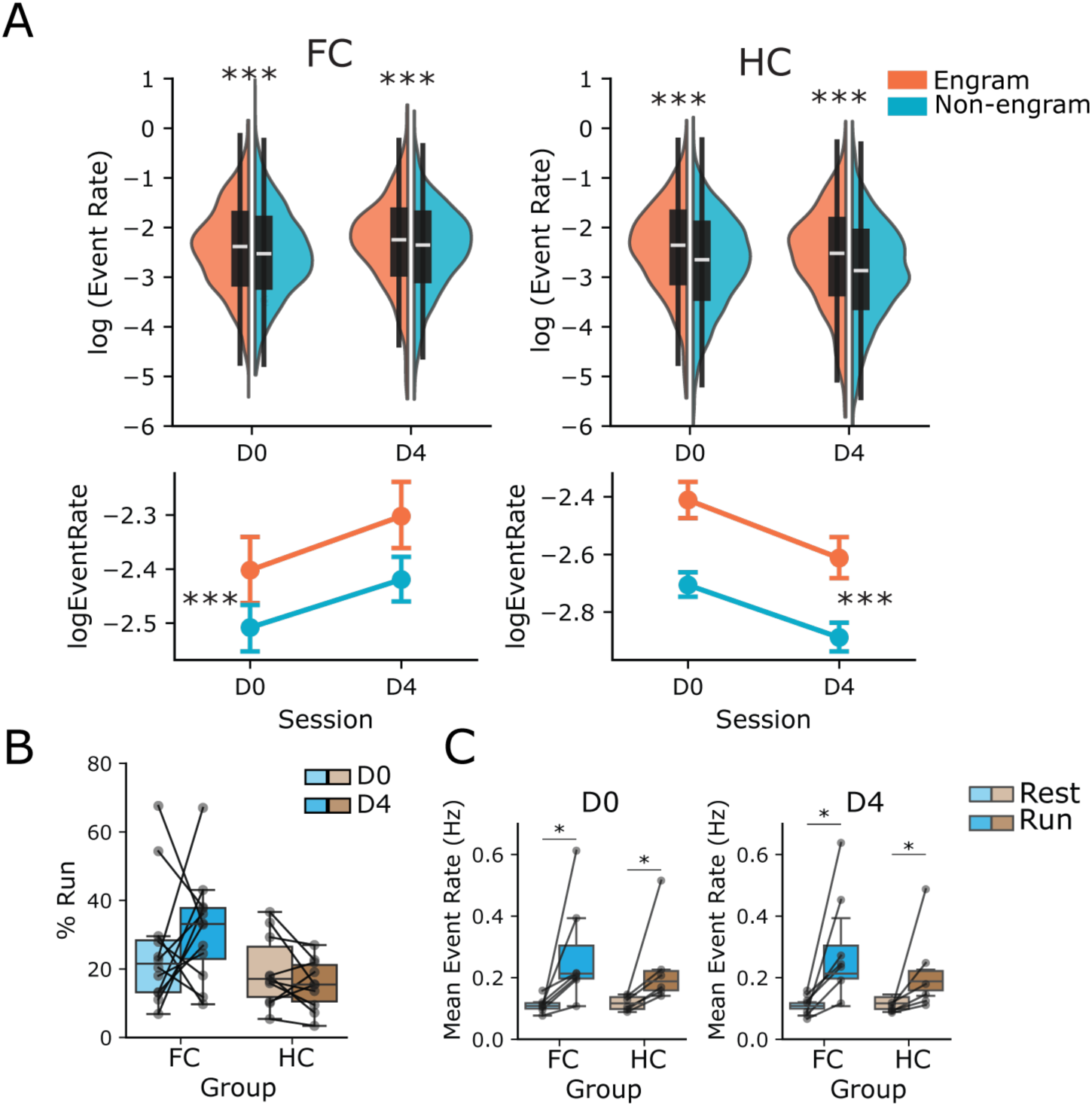
Effect of locomotion on event rates. A. Average calcium event rates across sessions for FC mice (n=8). Linear mixed effects model Group x Population x Session, Main Effect of Population *p=0.003,* Session *p=*0.143, Group *p*=0.354 Group x Session *p=*0.001. B. Average calcium event rates during rest periods across sessions for HC mice (n=8). Linear mixed effects model, Group x Population x Session, Main Effect of Population *p=0.003,* Session *p=*0.143, Group *p*=0.354 Group x Session p=0.001. C. % Time running for FC (blue) and HC (brown) groups. Mixed Effects Model, Group x Session, Group: *p=*0.3264, Session: *p=*0.2710, Interaction: *p=*0.2207. Points represent FOVs from FC (n=12 FOVs) or HC (n=10 FOVs) across mice. D. Mean event rates during rest and run periods on each Day. Run periods had significantly higher event rates. Left D0: Group p=0.9188, State, p=0.0039, Interaction, p=0.5758 Pairwise tests (Rest vs Run) FC: p=0.03125, HC p=0.0.03125. Right D4: Group, p=0.8557, p=0.0006, Interaction : p=0.3458. Pairwise tests (Rest vs Run) FC: p=0.03125, HC p=0.03125. Points represent animals from FC (n=7) or HC (n=6).

**Supplementary Figure 6.**
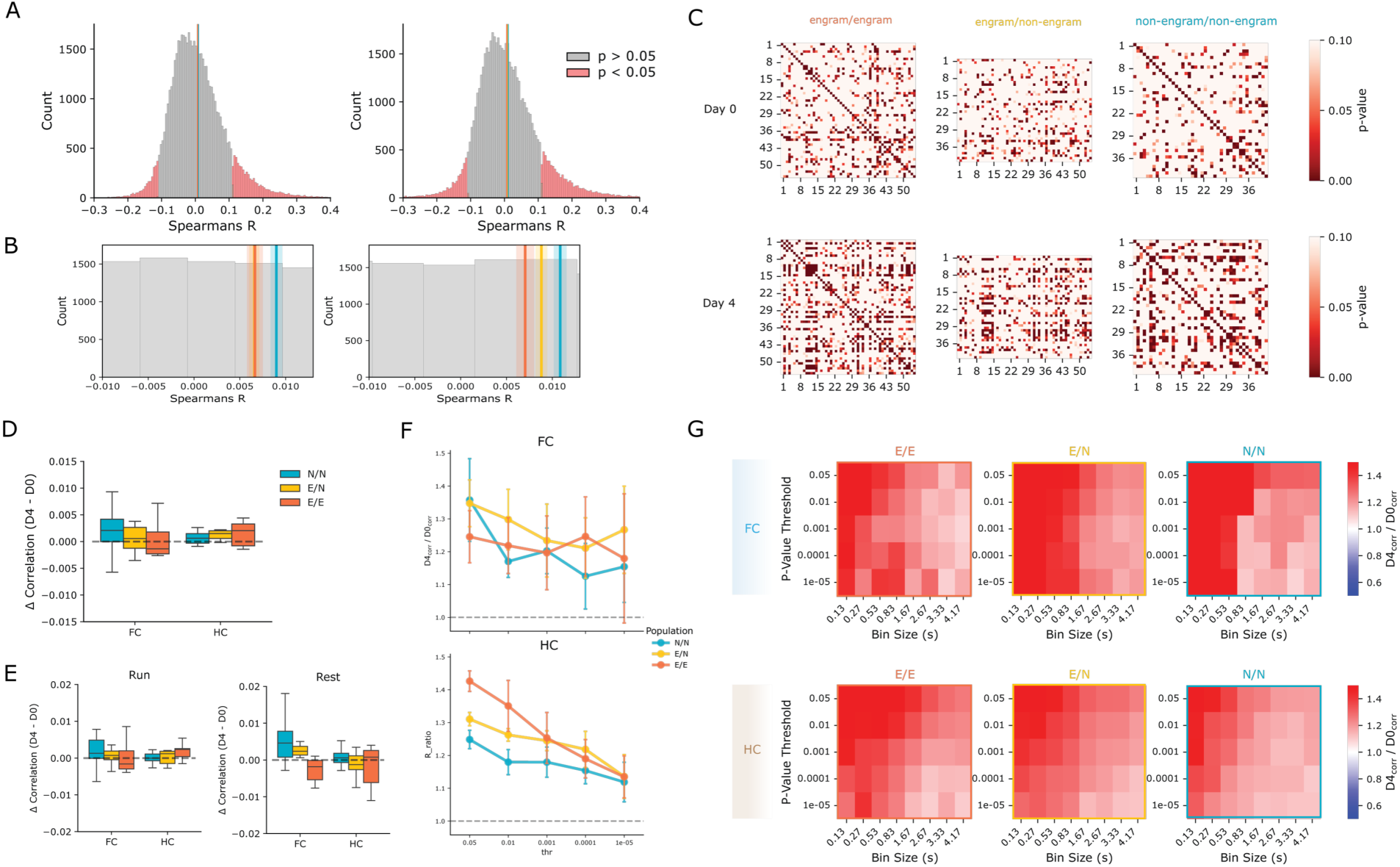
Additional correlation analyses. A) Histograms of all Spearman coefficient values with below insets of mean values across mice. Red areas indicate significantly correlated pairs analyzed in <u>Figures 4D-G</u>, and D-G. B) Inset of mean Spearmans R coefficient overlaid over distributions. The signifcantly correlated pairs exist in the tails of the distribution, rather than the population mean which is close to zero. C) P-values for example correlation matrix. A subsets of cells have significantly correlated coefficients. Correlations for <u>Figures 4C-D</u> were thresholded based on this p-value significance (p<0.05 to p<10e-5). D) No change in mean Spearman’s correlation across the whole session from D0 to D4 for pairwise correlations between engram/engram cells (orange: E/E), engram/non-engram cells (yellow: E-N), and non-engram/non-engram cells (teal: N/N). Linear mixed effects model, Group x Population. Main effect of Group *(p=*0.31), Population (*p=*0.53). A trend-level interaction was found for the FC x E/E effect compared to HC in N/N (*p=*0.07), consistent with increased running on D4. FC, N=8 mice HC, N=8 mice. E) No change in mean Spearman’s correlation during rest and run states after controlling for time. Left: Rest; Mixed ANOVA Group x Population, Main effect of Group: *p=*0.17, Population: *p=*0.18, Interaction: *p=*0.66. Right: Run; Linear mixed effects model Group x Population. Main effect of Group: *p=*0.45, Population: E/N *p=*0.93, E/E *p=*0.44, FC x E/E *p=*0.12. FC, N=7 mice HC, N=6 mice. F) Normalized change in correlations for thresholded cell pairs across significance levels. A bin size of 1.7 seconds was used. Top, FC Linear Mixed Effects Group X Population interaction. E/E X FC and E/N x FC interactions p=0.65 and p=0.68 respectively. Bottom HC Linear Mixed Effects Model Group X Population interaction. E/E X FC and E/N x FC interactions p=0.65 and p=0.68 respectively. FC, N=7 mice HC, N=6 mice. G) Heatmap of normalized change in correlation for thresholded pairs across timescales (bin sizes) and significance thresholds (p-values) in FC (top) mice and HC mice (bottom). Red: increase in correlation across days, Blue: decrease across days, white: no change. FC, N=7 mice HC, N=6 mice.

**Supplementary Figure 7.**
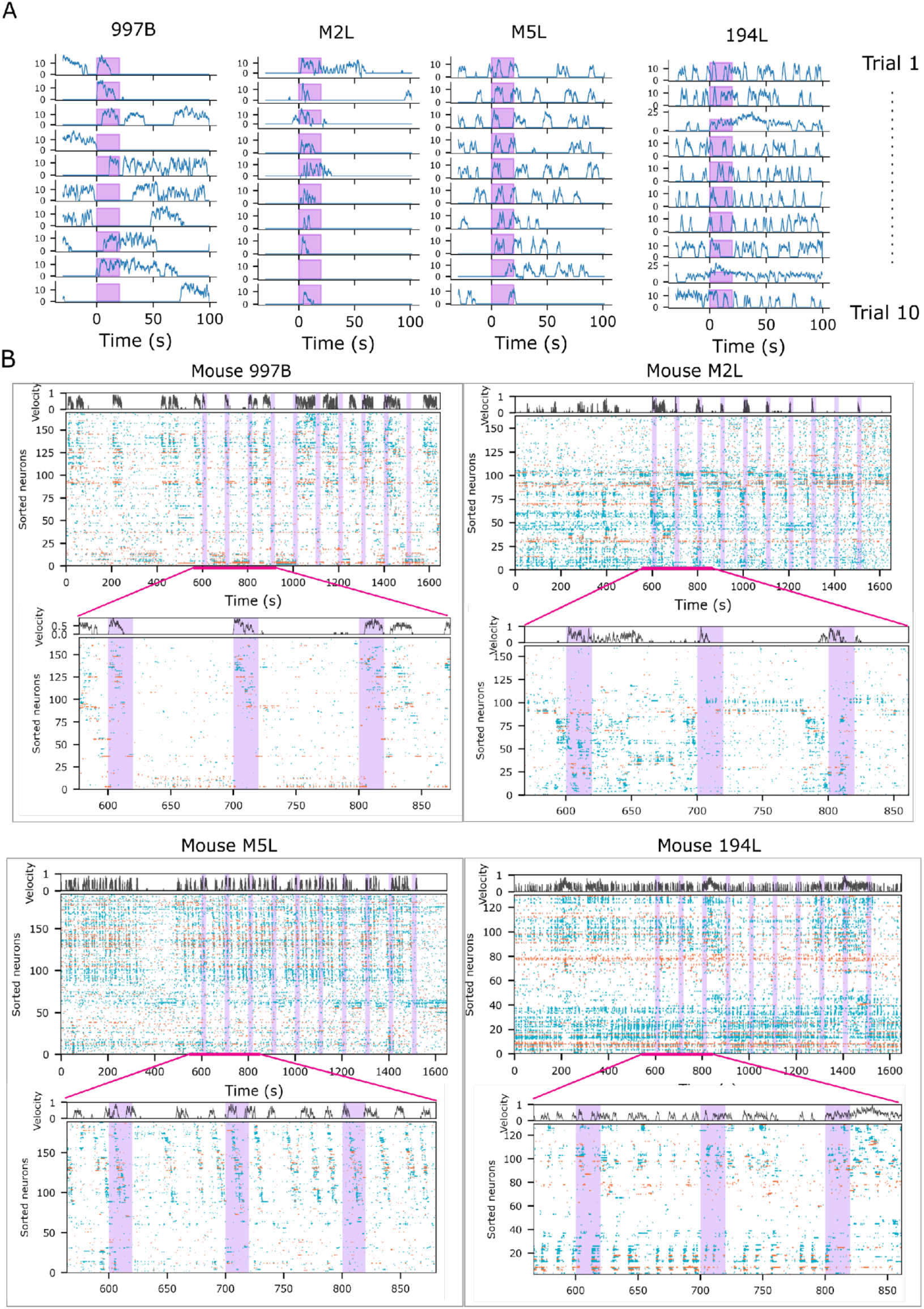
Tone Recall Rates & Locomotion across trials. A) 4 example mice running behavior across trials. Trials 1-10 are shown from top to bottom, and tone period is shown in purple. B) Additional rastermap-sorted raster plots for example mice (n=4). Beneath each raster is a zoomed inset of the first 3 trials (pink lines). Top: normalized velocity trace. Orange: engram cell calcium events, teal: non-engram cell calcium events. Purple: CS presentation

**Supplementary Figure 8.**
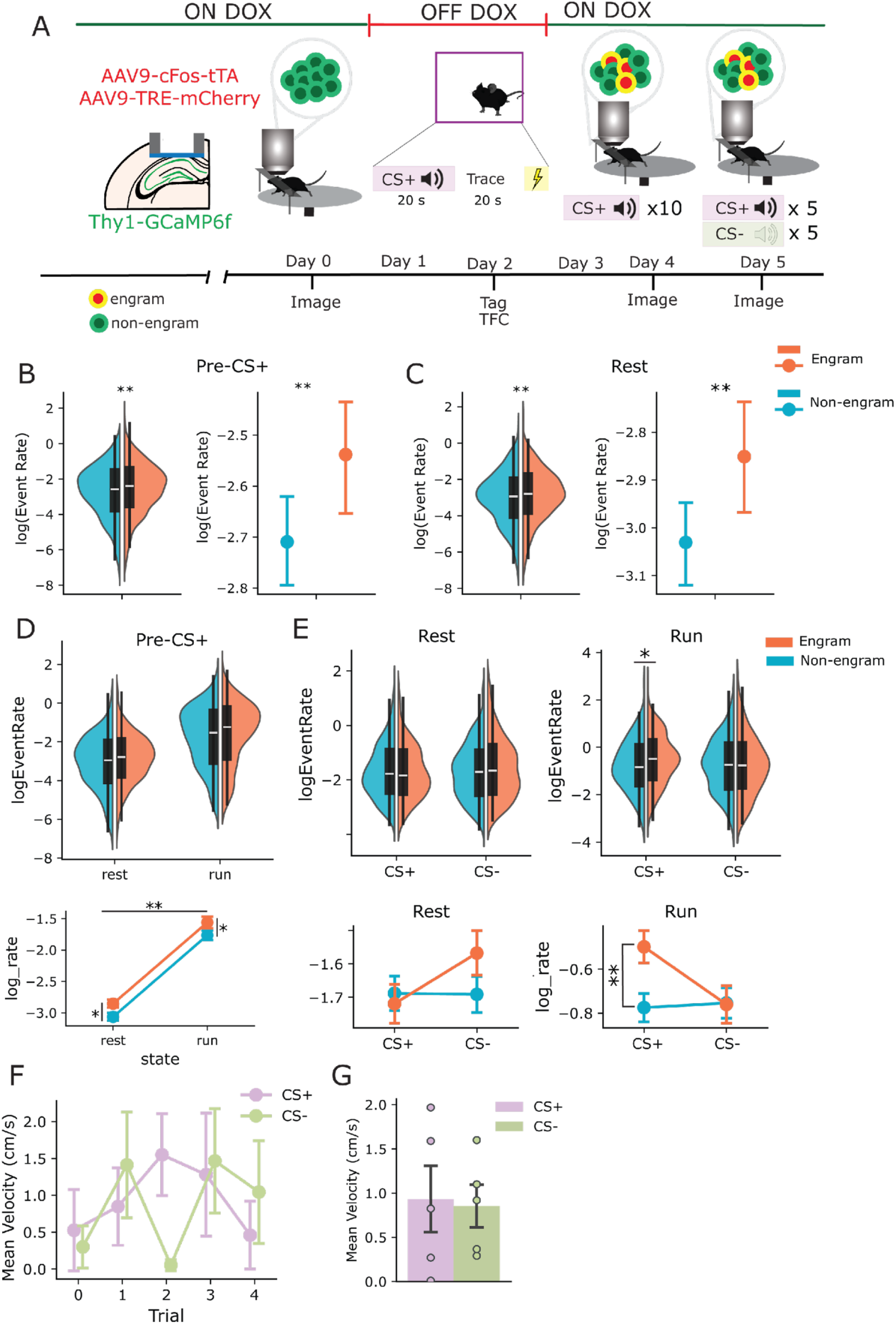
Neutral tone control experiment and additional rate analyses. A) Tone experimental timeline with additional day of neutral (CS-) and conditioned (CS+) tone presentation during imaging for n=5 mice on Day 5. CS+ and CS-were randomly interleaved (5 trials each) B) Engram cell rates during Day 4, before CS+ presentation. Linear Mixed Effects Model. Effect of Population *p=*0.005. Error bars represent 95% CI. C) Engram cell rates during Day 4, before CS+ presentation, Rest only. Linear Mixed Effects Model, Effect of Population *p=*0.004. Error bars represent 95% CI. D) Event rates during pre-CS+ baseline period. Linear Mixed Effects Model, Main Effect of Population *p=*0.03, Run *p=*2e-45. E) Rest versus Run event rates split by CS+/CS-trial types; Run activity is significantly higher in engram cells for the CS+. Linear Mixed Effects Model, Population x CS x Run, Main effect of Run *p=*1e-26, Engram x Run interaction *p=*0.016. Post hoc Wald test engram vs non-engram during Run *p=*0.0024 (bottom). F) Velocity across trace periods for CS+ and CS-trials. RM ANOVA, Effect of Trial *p=*0.5, CS *p=*0.86 G) Average trace period velocity for each trial type (paired t-test, *p=*0.86)

**Supplementary Figure 9.**
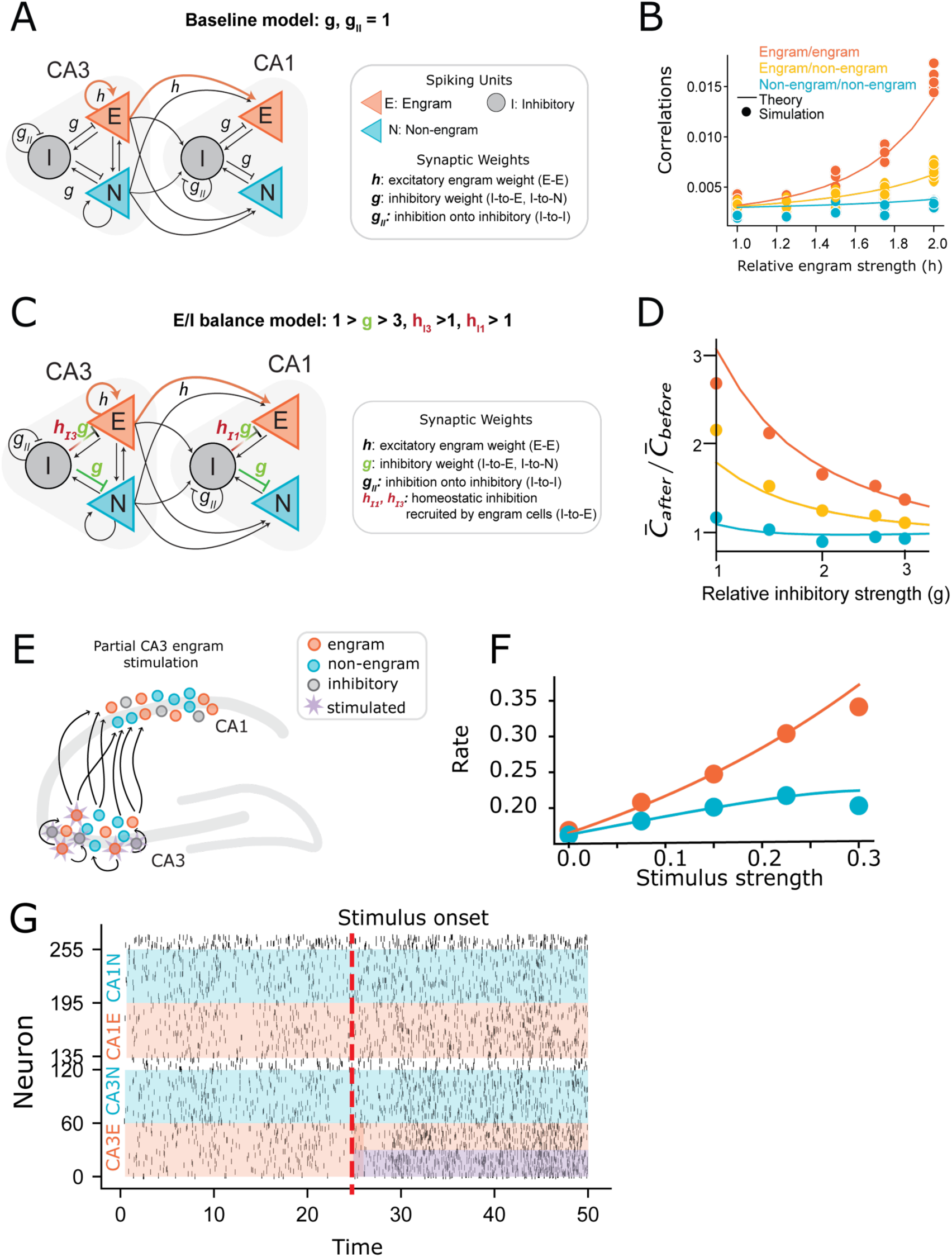
Engram Model Simulations. A) Schematic representation of model. A subset of cells are engram cells with excitatory weights specified by *h*. Inhibitory connections are specified by *g*, and inhibitory weights onto engram cells are scaled with *h_I3_ or h_I1_* for CA3 and CA1 respectively. B) In our baseline model, where inhibition and excitation have the same relative strength, correlations of engram cells (orange lines) increase a large amount as a function of engram strength *h*. C) Schematic representation of model with increased inhibition (E/I balance model). We introduced new parameters for homeostatic inhibition, hI3 and hI1, onto engram cells in addition to stronger global, non-specific inhibition (g > 1). D) Introducing homeostatic and global inhibition (strong E/I balance) dampens correlations in engram population. The ratio of correlations after potentiation of engram weights (*h* = 2) to before (*h* = 1) as a function of baseline inhibition *g*. E) Cartoon of stimulation procedure. A subset of excitatory Engram cells in CA3 received stimulation. F) Stimulating a portion of the CA3 engram cells paired with disinhibition (an inhibitory input to the CA1, CA3 inhibitory cells) leads to an increase in both CA1 engram rates and non-engram rates. G) Raster plot to test engram reactivation in the model. A subpopulation of CA3 engram cells received stimulation (time 25) reactivation.

## References

1. Josselyn, S.A., and Tonegawa, S. (2020). Memory engrams: Recalling the past and imagining the future. Science 367. 10.1126/science.aaw4325.

2. Josselyn, S.A., Köhler, S., and Frankland, P.W. (2015). Finding the engram. Nat Rev Neurosci 16, 521–534. 10.1038/nrn4000.

3. Eichenbaum, H. (2017). On the Integration of Space, Time, and Memory. Neuron 95, 1007– 1018. 10.1016/j.neuron.2017.06.036.

4. Denny, C.A., Kheirbek, M.A., Alba, E.L., Tanaka, K.F., Brachman, R.A., Laughman, K.B., Tomm, N.K., Turi, G.F., Losonczy, A., and Hen, R. (2014). Hippocampal memory traces are differentially modulated by experience, time, and adult neurogenesis. Neuron 83, 189–201. 10.1016/j.neuron.2014.05.018.

5. Liu, X., Ramirez, S., Pang, P.T., Puryear, C.B., Govindarajan, A., Deisseroth, K., and Tonegawa, S. (2012). Optogenetic stimulation of a hippocampal engram activates fear memory recall. Nature 484, 381–385. 10.1038/nature11028.

6. Ramirez, S., Liu, X., Lin, P.-A., Suh, J., Pignatelli, M., Redondo, R.L., Ryan, T.J., and Tonegawa, S. (2013). Creating a False Memory in the Hippocampus. Science 341, 387–391. 10.1126/science.1239073.

7. Shpokayte, M., McKissick, O., Guan, X., Yuan, B., Rahsepar, B., Fernandez, F.R., Ruesch, E., Grella, S.L., White, J.A., Liu, X.S., et al. (2022). Hippocampal cells segregate positive and negative engrams. Commun Biol 5, 1009. 10.1038/s42003-022-03906-8.

8. Chen, B.K., Murawski, N.J., Cincotta, C., McKissick, O., Finkelstein, A., Hamidi, A.B., Merfeld, E., Doucette, E., Grella, S.L., Shpokayte, M., et al. (2019). Artificially Enhancing and Suppressing Hippocampus-Mediated Memories. Current Biology, 1–10. 10.1016/j.cub.2019.04.065.

9. Grella, S.L., Fortin, A.H., Ruesch, E., Bladon, J.H., Reynolds, L.F., Gross, A., Shpokayte, M., Cincotta, C., Zaki, Y., and Ramirez, S. (2022). Reactivating hippocampal-mediated memories during reconsolidation to disrupt fear. Nat Commun 13, 4733. 10.1038/s41467-022-32246-8.

10. Zhou, Y., Zhu, H., Liu, Z., Chen, X., Su, X., Ma, C., Tian, Z., Huang, B., Yan, E., Liu, X., et al. (2019). A ventral CA1 to nucleus accumbens core engram circuit mediates conditioned place preference for cocaine. Nat Neurosci 22, 1986–1999. 10.1038/s41593-019-0524-y.

11. Tanaka, K.Z., He, H., Tomar, A., Niisato, K., Huang, A.J.Y., and McHugh, T.J. (2018). The hippocampal engram maps experience but not place. Science 361, 392–397. 10.1126/science.aat5397.

12. Pettit, N.L., Yap, E.-L., Greenberg, M.E., and Harvey, C.D. (2022). Fos ensembles encode and shape stable spatial maps in the hippocampus. Nature. 10.1038/s41586-022-05113-1.

13. Josselyn, S.A., and Frankland, P.W. (2018). Memory Allocation: Mechanisms and Function. Annual Review of Neuroscience 41, 389–413. 10.1146/annurev-neuro-080317-061956.

14. Chen, L., Cummings, K.A., Mau, W., Zaki, Y., Dong, Z., Rabinowitz, S., Clem, R.L., Shuman, T., and Cai, D.J. (2020). The role of intrinsic excitability in the evolution of memory: Significance in memory allocation, consolidation, and updating. Neurobiology of Learning and Memory 173, 107266. 10.1016/j.nlm.2020.107266.

15. Tanaka, K.Z., Pevzner, A., Hamidi, A.B., Nakazawa, Y., Graham, J., and Wiltgen, B.J. (2014). Cortical Representations Are Reinstated by the Hippocampus during Memory Retrieval. Neuron 84, 347–354. 10.1016/j.neuron.2014.09.037.

16. Ghandour, K., Ohkawa, N., Fung, C.C.A., Asai, H., Saitoh, Y., Takekawa, T., Okubo-Suzuki, R., Soya, S., Nishizono, H., Matsuo, M., et al. (2019). Orchestrated ensemble activities constitute a hippocampal memory engram. Nat Commun 10, 2637. 10.1038/s41467-019-10683-2.

17. Ryan, T.J., Roy, D.S., Pignatelli, M., Arons, A., and Tonegawa, S. (2015). Engram cells retain memory under retrograde amnesia. Science 348, 1007–1013. 10.1126/science.aaa5542.

18. Choi, J.H., Sim, S.E., Kim, J. il, Choi, D.I.I., Oh, J., Ye, S., Lee, J., Kim, T.H., Ko, H.G., Lim, C.S., et al. (2018). Interregional synaptic maps among engram cells underlie memory formation. Science 360, 430–435. 10.1126/science.aas9204.

19. Castello-Waldow, T.P., Weston, G., Ulivi, A.F., Chenani, A., Loewenstein, Y., Chen, A., and Attardo, A. (2020). Hippocampal neurons with stable excitatory connectivity become part of neuronal representations. PLoS Biol 18, e3000928. 10.1371/journal.pbio.3000928.

20. Uytiepo, M., Zhu, Y., Bushong, E., Chou, K., Polli, F.S., Zhao, E., Kim, K.-Y., Luu, D., Chang, L., Yang, D., et al. (2025). Synaptic architecture of a memory engram in the mouse hippocampus. Science 387, eado8316. 10.1126/science.ado8316.

21. Kim, S.J., Slocum, A.H., and Scott, B.B. (2022). A miniature kinematic coupling device for mouse head fixation. Journal of Neuroscience Methods 372, 109549. 10.1016/j.jneumeth.2022.109549.

22. Joo, H.R., and Frank, L.M. (2018). The hippocampal sharp wave–ripple in memory retrieval for immediate use and consolidation. Nat Rev Neurosci 19, 744–757. 10.1038/s41583-018-0077-1.

23. Carr, M.F., Jadhav, S.P., and Frank, L.M. (2011). Hippocampal replay in the awake state: a potential substrate for memory consolidation and retrieval. Nature neuroscience 14, 147–153.

24. Buzsáki, G., and Tingley, D. (2018). Space and Time: The Hippocampus as a Sequence Generator. Trends in Cognitive Sciences 22, 853–869. 10.1016/j.tics.2018.07.006.

25. Stringer, C., Zhong, L., Syeda, A., Du, F., Kesa, M., and Pachitariu, M. (2025). Rastermap: a discovery method for neural population recordings. Nat Neurosci 28, 201–212. 10.1038/s41593-024-01783-4.

26. Rao-Ruiz, P., Visser, E., Mitrić, M., Smit, A.B., and Van Den Oever, M.C. (2021). A Synaptic Framework for the Persistence of Memory Engrams. Front. Synaptic Neurosci. 13, 661476. 10.3389/fnsyn.2021.661476.

27. Gava, G.P., McHugh, S.B., Lefèvre, L., Lopes-dos-Santos, V., Trouche, S., El-Gaby, M., Schultz, S.R., and Dupret, D. (2021). Integrating new memories into the hippocampal network activity space. Nat Neurosci 24, 326–330. 10.1038/s41593-021-00804-w.

28. Venkatesh, M., Jaja, J., and Pessoa, L. (2020). Comparing functional connectivity matrices: A geometry-aware approach applied to participant identification. NeuroImage 207, 116398. 10.1016/j.neuroimage.2019.116398.

29. Yiu, A.P., Mercaldo, V., Yan, C., Richards, B., Rashid, A.J., Hsiang, H.-L.L., Pressey, J., Mahadevan, V., Tran, M.M., Kushner, S.A., et al. (2014). Neurons Are Recruited to a Memory Trace Based on Relative Neuronal Excitability Immediately before Training. Neuron 83, 722– 735. 10.1016/j.neuron.2014.07.017.

30. Rashid, A.J., Yan, C., Mercaldo, V., Hsiang, H.L., Park, S., Cole, C.J., De Cristofaro, A., Yu, J., Ramakrishnan, C., Lee, S.Y., et al. (2016). Competition between engrams influences fear memory formation and recall. Science 353. 10.1126/science.aaf0594.

31. Zhou, Y., Won, J., Karlsson, M.G., Zhou, M., Rogerson, T., Balaji, J., Neve, R., Poirazi, P., and Silva, A.J. (2009). CREB regulates excitability and the allocation of memory to subsets of neurons in the amygdala. Nat Neurosci 12, 1438–1443. 10.1038/nn.2405.

32. Park, S., Kramer, E.E., Mercaldo, V., Rashid, A.J., Insel, N., Frankland, P.W., and Josselyn, S.A. (2016). Neuronal Allocation to a Hippocampal Engram. Neuropsychopharmacology 41, 2987–2993. 10.1038/npp.2016.73.

33. Mocle, A.J., Ramsaran, A.I., Jacob, A.D., Rashid, A.J., Luchetti, A., Tran, L.M., Richards, B.A., Frankland, P.W., and Josselyn, S.A. (2024). Excitability mediates allocation of pre-configured ensembles to a hippocampal engram supporting contextual conditioned threat in mice. Neuron, S0896627324000916. 10.1016/j.neuron.2024.02.007.

34. Cai, D.J., Aharoni, D., Shuman, T., Shobe, J., Biane, J., Song, W., Wei, B., Veshkini, M., La-Vu, M., Lou, J., et al. (2016). A shared neural ensemble links distinct contextual memories encoded close in time. Nature 534, 115–118. 10.1038/nature17955.

35. Zaki, Y., Pennington, Z.T., Morales-Rodriguez, D., Bacon, M.E., Ko, B., Francisco, T.R., LaBanca, A.R., Sompolpong, P., Dong, Z., Lamsifer, S., et al. (2025). Offline ensemble co-reactivation links memories across days. Nature 637, 145–155. 10.1038/s41586-024-08168-4.

36. Chen, L., Francisco, T.R., Baggetta, A.M., Zaki, Y., Ramirez, S., Clem, R.L., Shuman, T., and Cai, D.J. (2023). Ensemble-specific deficit in neuronal intrinsic excitability in aged mice. Neurobiology of Aging 123, 92–97. 10.1016/j.neurobiolaging.2022.12.007.

37. Delamare, G., Tomé, D.F., and Clopath, C. (2024). Intrinsic neural excitability biases allocation and overlap of memory engrams. J. Neurosci., e0846232024. 10.1523/JNEUROSCI.0846-23.2024.

38. Crestani, A.P., Krueger, J.N., Barragan, E.V., Nakazawa, Y., Nemes, S.E., Quillfeldt, J.A., Gray, J.A., and Wiltgen, B.J. (2019). Metaplasticity contributes to memory formation in the hippocampus. Neuropsychopharmacology 44, 408–414. 10.1038/s41386-018-0096-7.

39. Mau, W., Hasselmo, M.E., and Cai, D.J. (2020). The brain in motion: How ensemble fluidity drives memory-updating and flexibility. eLife 9, e63550. 10.7554/eLife.63550.

40. Climer, J.R., Davoudi, H., Oh, J.Y., and Dombeck, D.A. (2025). Hippocampal representations drift in stable multisensory environments. Nature 645, 457–465. 10.1038/s41586-025-09245-y.

41. Dragoi, G., and Tonegawa, S. (2011). Preplay of future place cell sequences by hippocampal cellular assemblies. Nature 469, 397–401. 10.1038/nature09633.

42. Suthard, R.L., Senne, R.A., Buzharsky, M.D., Diep, A.H., Pyo, A.Y., and Ramirez, S. (2024). Engram reactivation mimics cellular signatures of fear. Cell Reports 43, 113850. 10.1016/j.celrep.2024.113850.

43. Cowansage, K.K., Shuman, T., Dillingham, B.C., Chang, A., Golshani, P., and Mayford, M. (2014). Direct Reactivation of a Coherent Neocortical Memory of Context. Neuron 84, 432–441. 10.1016/j.neuron.2014.09.022.

44. Javed, M.H., Robles-Hernandez, E.M., Patel, R., Haberl, M.G., and Da Silva, S.V. (2024). cfos principal cells and interneurons are strongly reactivated by sharp wave ripples. Preprint, 10.1101/2024.12.17.628897 https://doi.org/10.1101/2024.12.17.628897.

45. Jadhav, S.P., Kemere, C., German, P.W., and Frank, L.M. (2012). Awake Hippocampal Sharp-Wave Ripples Support Spatial Memory. Science 336, 1454–1458. 10.1126/science.1217230.

46. Villette, V., Malvache, A., Tressard, T., Dupuy, N., and Cossart, R. (2015). Internally Recurring Hippocampal Sequences as a Population Template of Spatiotemporal Information. Neuron 88, 357–366. 10.1016/j.neuron.2015.09.052.

47. Rajasethupathy, P., Sankaran, S., Marshel, J.H., Kim, C.K., Ferenczi, E., Lee, S.Y., Berndt, A., Ramakrishnan, C., Jaffe, A., Lo, M., et al. (2015). Projections from neocortex mediate top-down control of memory retrieval. Nature 526, 653–659. 10.1038/nature15389.

48. Tayler, K.K., Tanaka, K.Z., Reijmers, L.G., and Wiltgen, B.J. (2013). Reactivation of Neural Ensembles during the Retrieval of Recent and Remote Memory. Current Biology 23, 99–106. 10.1016/j.cub.2012.11.019.

49. Jou, C., Hurtado, J. R., Cruz-Holden, D., Carrillo-Segura, S., Park, E. H., and Fenton, A. A. (2025). Optogenetic stimulation of memory-tagged neurons elicits endogenous patterns of neural activity. bioRxiv. 10.1101/2023.05.15.540888.

50. Jung, J.H., Wang, Y., Mocle, A.J., Zhang, T., Köhler, S., Frankland, P.W., and Josselyn, S.A. (2023). Examining the engram encoding specificity hypothesis in mice. Neuron 111, 1830–1845.e5. 10.1016/j.neuron.2023.03.007.

51. Ahmed, M.S., Priestley, J.B., Castro, A., Stefanini, F., Solis Canales, A.S., Balough, E.M., Lavoie, E., Mazzucato, L., Fusi, S., and Losonczy, A. (2020). Hippocampal Network Reorganization Underlies the Formation of a Temporal Association Memory. Neuron 107, 283–291.e6. 10.1016/j.neuron.2020.04.013.

52. MacDonald, C.J., Carrow, S., Place, R., and Eichenbaum, H. (2013). Distinct hippocampal time cell sequences represent odor memories in immobilized rats. J Neurosci 33, 14607–14616. 10.1523/JNEUROSCI.1537-13.2013.

53. Taxidis, J., Pnevmatikakis, E.A., Dorian, C.C., Mylavarapu, A.L., Arora, J.S., Samadian, K.D., Hoffberg, E.A., and Golshani, P. (2020). Differential Emergence and Stability of Sensory and Temporal Representations in Context-Specific Hippocampal Sequences. Neuron 108, 984–998.e9. 10.1016/j.neuron.2020.08.028.

54. Ma, M., Simoes De Souza, F., Futia, G.L., Anderson, S.R., Riguero, J., Tollin, D., Gentile-Polese, A., Platt, J.P., Steinke, K., Hiratani, N., et al. (2024). Sequential activity of CA1 hippocampal cells constitutes a temporal memory map for associative learning in mice. Current Biology 34, 841–854.e4. 10.1016/j.cub.2024.01.021.

55. Wilmot, J.H., Diniz, C.R., Crestani, A.P., Puhger, K.R., Roshgadol, J., Tian, L., and Wiltgen, B.J. (2024). Phasic locus coeruleus activity enhances trace fear conditioning by increasing dopamine release in the hippocampus. eLife 12, RP91465. 10.7554/eLife.91465.

56. Puhger, K., Crestani, A.P., Diniz, C.R.A.F., and Wiltgen, B.J. (2024). The hippocampus contributes to retroactive stimulus associations during trace fear conditioning. iScience 27, 109035. 10.1016/j.isci.2024.109035.

57. Lucchetti, J., Fracasso, C., Balducci, C., Passoni, A., Forloni, G., Salmona, M., and Gobbi, M. (2019). Plasma and Brain Concentrations of Doxycycline after Single and Repeated Doses in Wild-Type and APP23 Mice. J Pharmacol Exp Ther 368, 32–40. 10.1124/jpet.118.252064.

58. Reijmers, L.G., Perkins, B.L., Matsuo, N., and Mayford, M. (2007). Localization of a stable neural correlate of associative memory. Science 317, 1230–1233. 10.1126/science.1143839.

59. DeNardo, L.A., Liu, C.D., Allen, W.E., Adams, E.L., Friedmann, D., Fu, L., Guenthner, C.J., Tessier-Lavigne, M., and Luo, L. (2019). Temporal evolution of cortical ensembles promoting remote memory retrieval. Nat Neurosci 22, 460–469. 10.1038/s41593-018-0318-7.

60. De Sousa, A.F., Zeidler, Z.E., Almeida-Filho, D.G., Shen, Y., Luchetti, A., Simanian, S., Mardini, M., DeNardo, L.A., and Silva, A.J. (2026). The prefrontal cortex controls memory organization in the hippocampus. Nat Neurosci 29, 1191–1202. 10.1038/s41593-026-02231-1.

61. DeNardo, L., and Luo, L. (2017). Genetic strategies to access activated neurons. Current Opinion in Neurobiology 45, 121–129. 10.1016/j.conb.2017.05.014.

62. Hyun, J.H., Nagahama, K., Namkung, H., Mignocchi, N., Roh, S.-E., Hannan, P., Krüssel, S., Kwak, C., McElroy, A., Liu, B., et al. (2022). Tagging active neurons by soma-targeted Cal-Light. Nat Commun 13, 7692. 10.1038/s41467-022-35406-y.

63. Sanchez, M.I., Nguyen, Q.-A., Wang, W., Soltesz, I., and Ting, A.Y. (2020). Transcriptional readout of neuronal activity via an engineered Ca ^2+^ -activated protease. Proc. Natl. Acad. Sci. U.S.A. 117, 33186–33196. 10.1073/pnas.2006521117.

64. Kim, C.K., Sanchez, M.I., Hoerbelt, P., Fenno, L.E., Malenka, R.C., Deisseroth, K., and Ting, A.Y. (2020). A Molecular Calcium Integrator Reveals a Striatal Cell Type Driving Aversion. Cell 183, 2003–2019.e16. 10.1016/j.cell.2020.11.015.

65. Jung, J.H., Wang, Y., Rashid, A.J., Zhang, T., Frankland, P.W., and Josselyn, S.A. (2023). Examining memory linking and generalization using scFLARE2, a temporally precise neuronal activity tagging system. Cell Reports 42, 113592. 10.1016/j.celrep.2023.113592.

66. Sheintuch, L., Rubin, A., Brande-Eilat, N., Geva, N., Sadeh, N., Pinchasof, O., and Ziv, Y. (2017). Tracking the Same Neurons across Multiple Days in Ca2+ Imaging Data. Cell Reports 21, 1102–1115. 10.1016/j.celrep.2017.10.013.

67. Giovannucci, A., Friedrich, J., Gunn, P., Kalfon, J., Brown, B.L., Koay, S.A., Taxidis, J., Najafi, F., Gauthier, J.L., Zhou, P., et al. (2019). CaImAn an open source tool for scalable calcium imaging data analysis. eLife 8, e38173. 10.7554/eLife.38173.

68. Stringer, C., Wang, T., Michaelos, M., and Pachitariu, M. (2021). Cellpose: a generalist algorithm for cellular segmentation. Nat Methods 18, 100–106. 10.1038/s41592-020-01018-x.

69. Dana, H., Chen, T.-W., Hu, A., Shields, B.C., Guo, C., Looger, L.L., Kim, D.S., and Svoboda, K. (2014). Thy1-GCaMP6 Transgenic Mice for Neuronal Population Imaging In Vivo. PLoS ONE 9, e108697. 10.1371/journal.pone.0108697.

70. Jewell, S., and Witten, D. (2018). Exact spike train inference via $\ell_{0}$ optimization. Ann. Appl. Stat. 12. 10.1214/18-AOAS1162.

71. Jewell, S.W., Hocking, T.D., Fearnhead, P., and Witten, D.M. (2020). Fast nonconvex deconvolution of calcium imaging data. Biostatistics 21, 709–726. 10.1093/biostatistics/kxy083.

72. De Vries, S.E.J., Lecoq, J.A., Buice, M.A., Groblewski, P.A., Ocker, G.K., Oliver, M., Feng, D., Cain, N., Ledochowitsch, P., Millman, D., et al. (2020). A large-scale standardized physiological survey reveals functional organization of the mouse visual cortex. Nat Neurosci 23, 138–151. 10.1038/s41593-019-0550-9.

73. Hawkes, A.G. (1971). Spectra of some self-exciting and mutually exciting point processes. Biometrika 58, 83–90. 10.1093/biomet/58.1.83.

74. Pernice, V., Staude, B., Cardanobile, S., and Rotter, S. (2011). How Structure Determines Correlations in Neuronal Networks. PLoS Comput Biol 7, e1002059. 10.1371/journal.pcbi.1002059.

75. Ocker, G.K., Josić, K., Shea-Brown, E., and Buice, M.A. (2017). Linking structure and activity in nonlinear spiking networks. PLoS Comput Biol 13, e1005583. 10.1371/journal.pcbi.1005583.

76. Ocker, G.K., and Doiron, B. (2019). Training and Spontaneous Reinforcement of Neuronal Assemblies by Spike Timing Plasticity. Cerebral Cortex 29, 937–951. 10.1093/cercor/bhy001.

